# Gene2Vec: Distributed Representation of Genes Based on Co-Expression

**DOI:** 10.1101/286096

**Authors:** Jingcheng Du, Peilin Jia, Yulin Dai, Cui Tao, Zhongming Zhao, Degui Zhi

**Author notes:** Corresponding Author: Degui Zhi, Ph.D. equal contribution (Jingcheng Du, Peilin Jia). equal contribution (Zhongming Zhao, Degui Zhi).

## Abstract

**Background.:** Existing functional description of genes are categorical, discrete, and mostly through manual process. In this work, we explore the idea of gene embedding, distributed representation of genes, in the spirit of word embedding.

**Methods & Results.:** From a pure data-driven fashion, we trained a 200-dimension vector representation of all human genes, using gene co-expression patterns in 984 data sets from the GEO databases. These vectors capture functional relatedness of genes in terms of recovering known pathways - the average inner product (similarity) of genes within a pathway is 1.52X greater than that of random genes. Using t-SNE, we produced a gene co-expression map that shows local concentrations of tissue specific genes. We also illustrated the usefulness of the embedded gene vectors, laden with rich information on gene co-expression patterns, in tasks such as gene-gene interaction prediction.

**Conclusions.:** We proposed a machine learning method that utilizes transcriptome-wide gene co-expression to generate a distributed representation of genes. We further demonstrated the utility of our distribution by predicting gene-gene interaction based solely on gene names. The distributed representation of genes could be useful for more bioinformatics applications.

## INTRODUCTION

Genes, discrete segments of the genome that are transcribed, are basic building blocks of molecular biological systems. Although almost all transcripts in the human genome have been identified, functional annotation of genes is still a challenging task. Most existing annotation efforts organize genes into functional categories, e.g., pathways, or represent their relationship into networks. Pathways and networks crystallize biological knowledge and are convenient qualitative conceptualization of gene functions. Yet the exact functions of a gene are often more subtle and elusive to be expressed in qualitative terms.

The challenge of creating a quantitative semantic representation of discrete units of a complex system is not unique to gene systems. For a long time, creating a quantitative representation of words had been challenging for linguistic modeling. Hinton proposed the pioneering idea of ‘learning distributed representations of words’ [1], i.e., representing the semantics of a word by mapping them to vectors in a high-dimension space. However, Hinton’s idea did not lead to real implementation in mainstream natural language processing (NLP) research, until recently. The word2vec model achieved success in NLP modeling [2]. This process of distributed representation is often called neural embedding because the embedding function is often expressed by a neural network with a large number of parameters. This success of word2vec inspires us to investigate the possibility to represent gene functions via neural embedding.

In this study, we aim to represent genes as vectors in a high-dimension space, i.e., a gene embedding. In the word2vec model [2], a word embedding is trained by maximizing the probability of word co-occurrences in context, i.e., only a few words apart in a same sentence. Analogously, we defined the context of a gene by the other genes that co-expressed with it. We derive an embedding such that the probability of the context of a gene is maximized. While it is possible to train a gene embedding using the standard NLP word embedding by using a biomedical corpus, such as the PubMed abstracts, published literature is incomplete and biased towards genes that are well-studied. Therefore, we intended to adopt a purely data-driven fashion.

Using co-expression patterns of all human genes in 984 whole transcriptome human gene expression data sets from Gene Expression Omnibus (GEO), we learned a gene embedding using a neural network. We show that our embedding grouped related genes in clusters. Moreover, we demonstrate the usefulness of the learned gene embedding to downstream tasks in the problem of prediction of gene-gene interaction.

## METHODS

### Data collection

#### Overview

We chose to use GEO data with rationale from both biological and technical aspects. In cellular systems, the mRNA expression levels represent activities of genes with fine resolutions. Over the past 10–20 years, GEO deposits the majority of microarray-based gene expression data in various conditions. Although the recent development of RNA-sequencing has generated transcriptomic data with advantages in both accuracy and scales than array-based data, the large cohort of GEO data provides features that are more suitable for our work. GEO data have been curated for over ten years and hence, the measurement covers a wide range of cell and tissue types, cellular conditions, disease status, and developmental stages. As our ultimate goal is to build a gene co-expression map that could be used for inferences in various conditions, we collected GEO data for our task. In addition, we chose one single platform to reduce technical variability and required the organism to be Homo sapiens.

#### Gene expression

We used the keywords “expression and human” to search in GEO on 12/24/2017 and retrieved all GSE sets that were conducted using the platform Affymetrix Human Genome U133 Plus 2.0 Array (GPL570). We required each dataset to have ≥ 30 samples. The downloaded gene expression intensity data were log transformed and quantile-normalized. For genes with multiple probe sets, we chose the probe set with the largest variance across all samples. Gene co-expression was measured using Pearson Correlation Coefficient (PCC) for each data set. In each data set, gene pairs with the PCC ≥ 0.9 were selected for following analysis. Selected gene pairs from all data sets were merged and serve as training data. We did not distinguish biological conditions.

#### Gene types on chip

The U133 array is one of the most widely utilized platform to measure human gene expression. The chip has 54,675 probe sets for 24,442 genes. The number of probe sets per gene ranged between 1 and 15, with more than half of genes (52.08%) have one probe sets. Among these genes, 21,960 (89.85%) genes could be mapped to the current version of NCBI Entrez gene annotation. These mappable genes include 18,055 protein-coding genes, 2,660 ncRNA, 730 pseudo genes, 132 snoRNA, and 383 other types of genes. Particularly for ncRNAs, there are 202 microRNA genes. Gene set enrichment analysis was conducted using Fisher’s Exact Test.

#### Gene-gene interaction dataset

We followed previous work [3] to build datasets for genegene interaction based on shared Gene Ontology (GO) annotations. GO annotation was obtained using the R (x64 3.4.3) package “org.Hs.eg.db” (version 3.5.0). GO structure file in the obo format was downloaded from [4]. All genes were mapped to NCBI Entrez Gene [5] (downloaded on 11/6/2017). We defined gene pairs that shared GO annotations as the positive set of functional association. To this end, we chose the GO category “Biological Process” with experimental evidence: IDA (inferred from direct assay), IMP (inferred from mutant phenotype), IPI (inferred from protein interaction), IGI (inferred from genetic interaction), and TAS (traceable author statement). To minimize generalized annotation, we excluded the highly over-represented GO terms including (1) “signal transduction” (GO:0007165); (2) three phosphorylation terms: “protein amino acid phosphorylation” (GO:0006468), “protein amino acid autophosphorylation” (GO:0046777), and “protein amino acid dephosphorylation” (GO:0006470); and (3) all terms at the first three levels of GO hierarchy (assuming the root term of biological process, “GO:0008150”, is level 0). This lead to a total of 270,704 pairs involving 5,369 genes. To build the negative data set, we obtained all gene-pairs that did not share any GO term or their children GO terms. This resulted a total of 40,879,714 gene pairs involved in 12,521 (64.85% of 19,307) human genes, serving as the set in which pairs of genes are not functionally associated.

#### Tissue-specific genes

GTEx data (version 6) [6] was used to estimate the tissue-specific expression pattern of genes in 27 tissues, each with ≥ 30 samples. For each gene, a z-score was calculated to measure its tissue specificity by comparing the average gene expression of the gene across all tissues [7].

#### Functional gene sets

We use clusteredness of MSigDB pathways (v5.1) [8] as the target function for hyper-parameter tuning for gene embedding training. Specifically, we used the category c2 including curated pathways from various online resources such as KEGG [9], Biocarta [10], and Reactome [11]. A total of 4,726 pathways were downloaded.

### Concept embedding of genes

Distributed representation of word, or neural word embedding, was a recent breakthrough in NLP research based on deep learning. The goal of word embedding is to derive a linear mapping, i.e., embedding, from the discrete space of individual words to a continuous Euclidean space such that similar words will be mapped to points in close vicinity in the embedding space. The direct benefit of word embedding is that such representation of individual words, vectors in continuous space, becomes differentiable and thus amenable for back-propagation-based neural network modeling. Meanwhile, a nice surprising result is that embedded space admits basic geometry. E.g., the KING - QUEEN ≈ MAN - WOMAN.

Inspired by the success of word embedding, we intend to produce an embedding of genes, also a discrete conceptual unit, such that similar genes are mapped to similar vectors. While for genes we do not have a natural equivalent concept of sentence in natural languages, we will use the notion of co-expression. This is analogy of the concept of co-occurrence in natural languages.

For neural embedding, a neural network is designed that maximizes an objective function, often in a form of likelihood, such as the probability of a word given its context. The most commonly used architectures are skip-gram and continuous bag-of-words (CBOW) that discussed in the word2vec approach [2]. In both architectures, a two-layer neural network is constructed to predict word co-occurrence, or the co-occurrence of a word and its surrounding words, or context. In CBOW, the input is the context and the output is the word; in skip-gram, the input is the word and the output is the context. For both architectures, input and output are connected through a middle projection layer. Note that neither neural network would offer satisfactory predictions for most of the words. But the real goal of word embedding is to learn a distributional representation, i.e., the parameters of the embedding mapping from the input to the middle projection layer. A simple fully connected linear layer was used for the embedding mapping. For CBOW, the embedded vectors of all words in the context are averaged and thus provide a uniform size vector for the next layer. The second layer for both architecture is a linear layer with a soft-max. A cross-entropy loss is minimized.

In gene embedding, we are using the genes who are co-expressed with the gene of interest as its context. Since the number of co-expressed genes may vary, the size of the context may vary as well. For simplicity, in this work, we extract all pairs of co-expressed genes and maximize the probability of one given the other for each pair. This is equivalent to the skipgram model. Since we are optimizing the total probability of all edges in a co-expression network, our approach can also be viewed as a graph embedding [12].

More formally, the input of the training problem is a list of gene pairs that are highly coexpressed, *T =* {(*g_i_*_1_*, g_i_*_2_)}, we will train an embedding network. The input of the network is a one-hot encoded vector for gene *g_i_* ∈ *R^d^*, where *d* is the number of genes and the elements of *g_i_* are all 0 expert *g_i_*[*x_i_*] *=* 1, where *x_i_* is the dimension corresponds to the gene *g_i_*. The output of the network is a vector of dimension *v_i_* ∈ *R^k^*, the embedding dimension. The parameter of the network is a matrix *W* ∈ *R^d×k^* such that 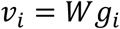. If we define the probability

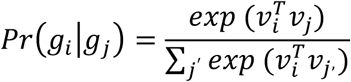

The loss function that is to be minimized is the negative likelihood 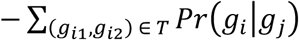. It can be shown that this complex loss function for this single layer network is equivalent to a two-layer network with shared weight matrices of *W* and *W^t^*, and the loss function as the standard cross-entropy after softmax (see, **Figure 1**).

**Figure 1.**
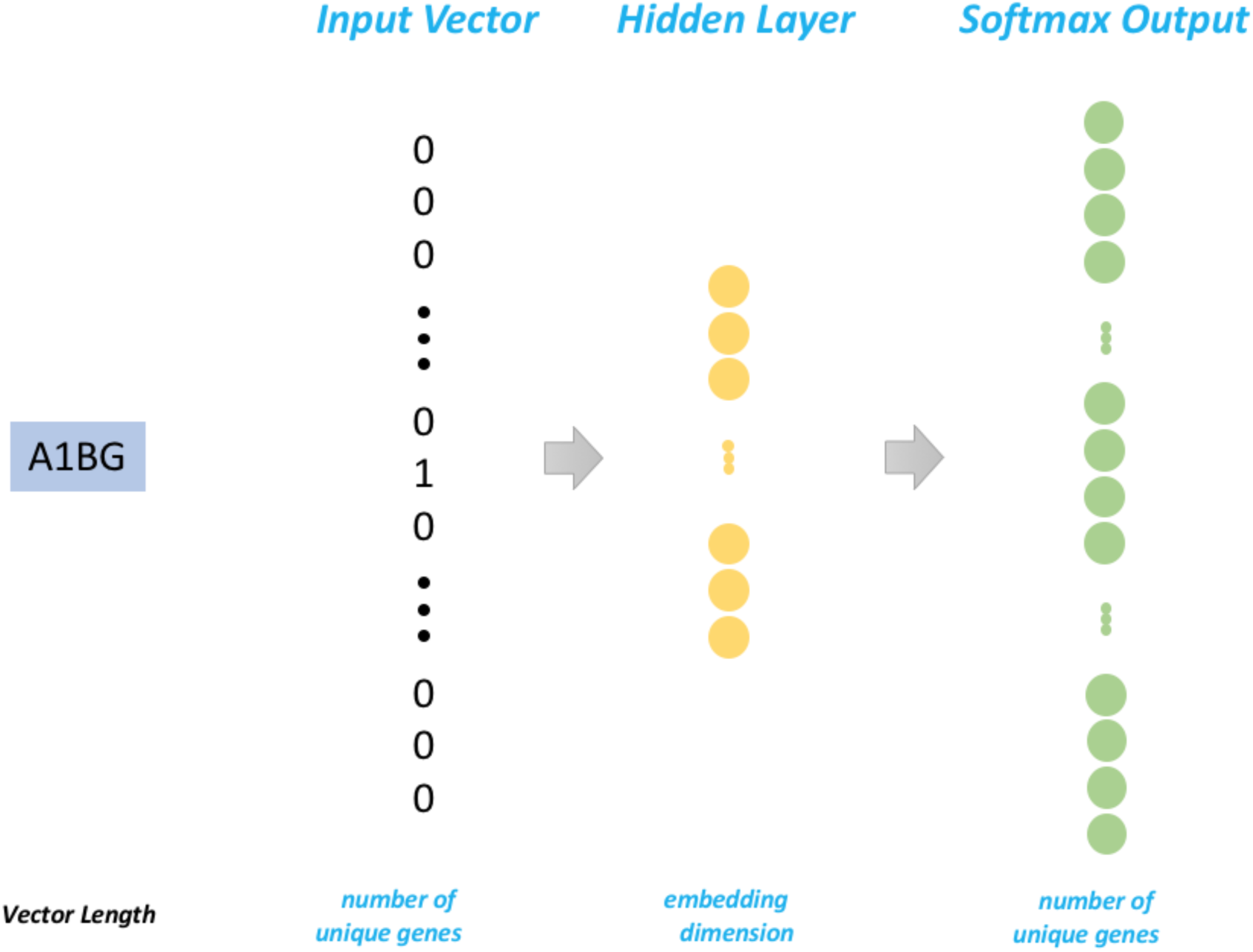
The Skip-Gram architecture was used for training for gene embedding. This is the modified architecture which is equivalent to the original word2vec, adopted from this blog [22]

### Training of embedding

We took all the gene pairs that have a PCC equal to or larger than 0.9 as the input. This is a choice due to limited computational resources. We shuffled the gene pairs in each dataset on every iteration to avoid the impact caused by the order of gene pairs in the datasets. The embedding was trained on all genes with a minimum frequency at 5. As number of iterations and dimensionality of the embedding are considered as two major hyper-parameters parameters for word embedding [13], in order to generate “best” gene embedding, we did a preliminary parameters tuning and performed a grid search to find best parameters. The search ranges for number of iterations and embedding dimension are set at 1 to 10 and 50, 100, 200, and 300 respectively. We used the word2vec function implemented in the gensim library [14] to generate gene embedding. Other parameters were set as default.

Since our goal is to obtain a gene embedding that reflects the functional relationships among genes, we selected the set of hyper-parameters that maximizes the clusteredness of genes within functional pathways. We optimized the following target function:

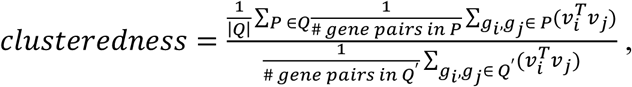

where *Q* is the set of pathways in MSigDB, and *Q’* is a set of random gene pairs. Due to the limitation of computation power, we selected all the pathways from the MSigDB with the number of genes equal or fewer than 50. In total, 6,729 pathways were selected as *Q*. We randomly selected 1,000 genes from gene embedding and generated all possible unique gene pairs (499,500 in total) as *Q’*.

### Visualization by t-SNE

A common way to visualize high-dimensional datasets is to map the datasets into 2D or 3D array. *t*-Distributed Stochastic Neighbor Embedding (t-SNE) is a machine learning algorithm for dimensionality reduction, which optimizes for neighborhood preserving and thus particularly well suited for the visualization of high-dimensional datasets [15]. Visualizations produced by t-SNE have been found significantly better than those produced by the other techniques [15].

In order to speed up the t-SNE on the high-dimensional gene embedding, we first reduced the dimension to 50 using PCA and then applied a multicore modification of Barnes-Hut t-SNE by L. Van der Maaten [16,17]. The perplexity was set at 30 and the learning rate was set at 200. To get stable t-SNE results, we set the number of iterations at 100,000.

### Prediction of gene-gene interaction

To investigate the usefulness of the trained gene embedding for downstream tasks, we applied the embedding to the problem of gene-gene interaction prediction. The goal is, given a pair of genes, we design a gene-gene interaction predictor neural network (GGIPNN) to predict if they will be together in any of the annotated pathway [3].

The architecture of GGIPNN can be seen in **Figure 2**. We first convert the genes in each gene pair to one-hot vectors and then map the one-hot vectors to gene embedding vectors using a shared embedding matrix. Then, the two gene embedding vectors will be concatenated together and be fed to a fully connected layer with a dimension at 100. The output will be fed to another fully connected layer with a dimension at 100. The output of the second fully connected layer will then be fed to a second fully connected layer with a dimension at 10. The output will be then fed to a softmax (same as sigmoid as this is a binary classification) layer. We compute the cross entropy of the softmax function output and then compute the mean of elements across results as the loss. We choose ReLU (Rectified Linear Units) as the activation function. To avoid overfitting, we apply dropout on both the first and second fully connected layers. The dropout out rates are set at 0.5.

**Figure 2.**
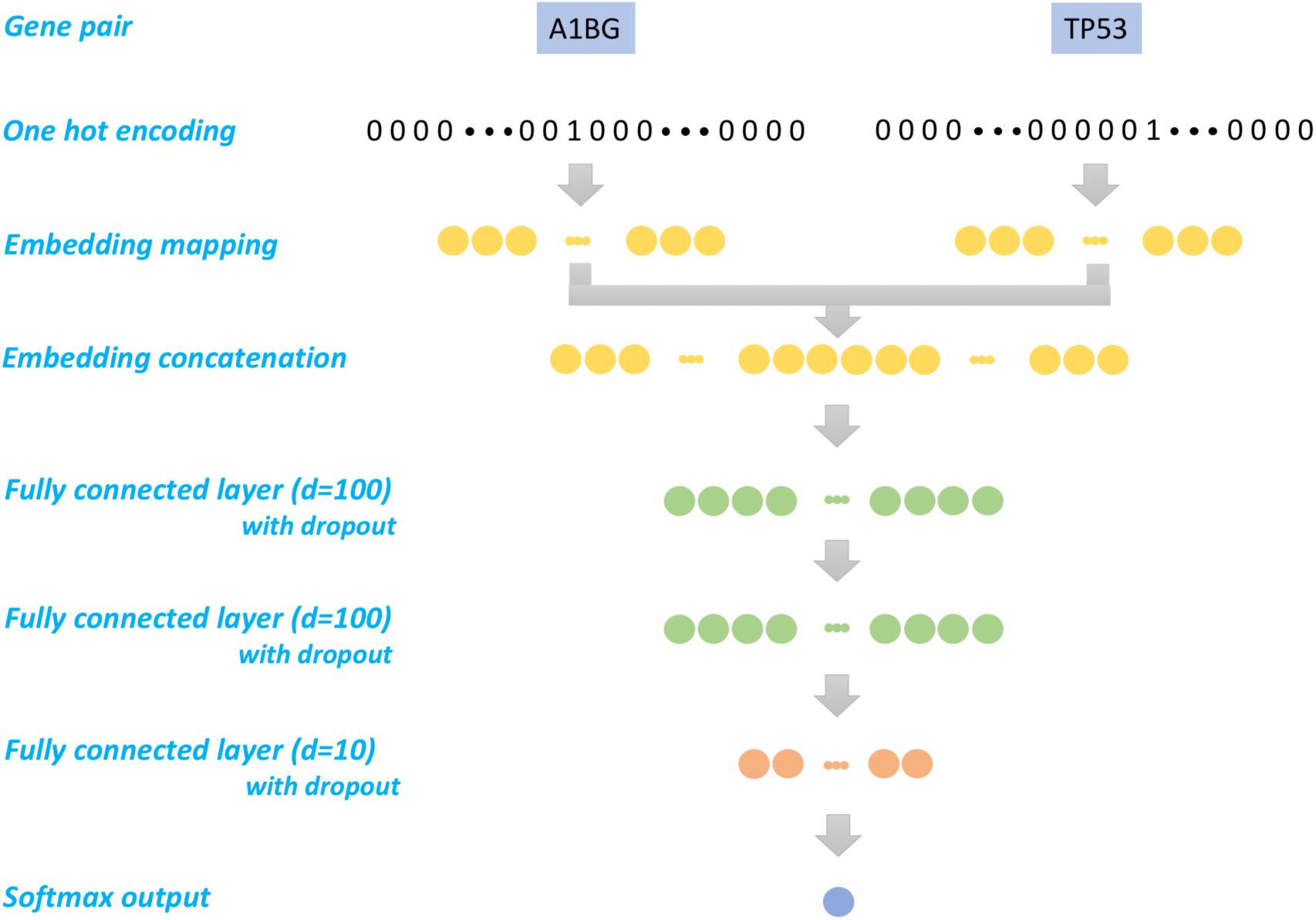
The architecture of gene-gene interaction predictor neural network (GGIPNN)

Area Under Curve (AUC) is computed to measure the performance of the prediction. We compared the AUC score of our pre-trained gene embedding and embedding which is randomly initialized. We also investigated the impact of trainable embedding layer (fine-tuning during the training) versus non-trainable embedding layer (fixed during the training) on the prediction. This model was implemented in TensorFlow.

We took all the pairs from HumanNet.v1.benchmark [3] as the positive pairs. This led to a total of 270,704 pairs involving 5,369 genes. To build the negative data set, we obtained all gene-pairs that did not share any GO term or their children GO terms. This resulted in a total of 40,879,714 gene-pairs involved in 12,521 (64.85% of 19,307) human genes, serving as the set in which pairs of genes are not functionally associated. To avoid the impact of the imbalanced labels distribution, we randomly selected negative pairs with the equal number of the positive pairs. We then split all the unique genes into training, validation and testing sets with an proportion of 7: 1: 2. The pairs that the both two genes belong to training set are used as training; the pairs that the both two genes belong to validation set are used as validation; the pairs that the both two genes belong to testing set are used as testing. By doing so, we avoid the possibility that the neural network “memorizes” the likelihood of genes to be interacting with any other genes. In total, the training dataset has 263,016 pairs (involving 8,832 genes), while the validation and testing dataset have 5,568 pairs (1,173 genes) and 21,448 pairs (2,467 genes) respectively.

## RESULTS

### Parameter tuning results by clusteredness

The parameter tuning results can be seen in Table 1. As we can observe, the dimension of 200 at iteration 9 produced best gene embedding using clusteredness as the target function (1.521). As result, we chose this embedding for all following analyses.

**Table 1.** Hyperparameter tuning using clusteredness as target function. **Bold** number denotes the largest number in that row.

| Dimension | Number of Iterations |  |  |  |  |  |  |  |  |  |
| --- | --- | --- | --- | --- | --- | --- | --- | --- | --- | --- |
|  | 1 | 2 | 3 | 4 | 5 | 6 | 7 | 8 | 9 | 10 |
| 50 | 1.428 | 1.444 | 1.467 | 1.470 | **1.487** | 1.465 | 1.473 | 1.479 | 1.475 | 1.462 |
| 100 | 1.415 | 1.467 | 1.488 | 1.491 | 1.498 | 1.501 | **1.519** | 1.486 | 1.480 | 1.490 |
| 200 | 1.403 | 1.463 | 1.491 | 1.498 | 1.495 | 1.482 | 1.470 | 1.488 | **1.521** | 1.509 |
| 300 | 1.392 | 1.443 | 1.472 | 1.473 | 1.473 | 1.509 | 1.474 | **1.513** | 1.479 | 1.480 |

### Gene embedding groups similar genes into spatial clusters

Using the first and second components from the t-SNE representation, we produced a gene co-expression map, based on which we explored the distribution of all human genes from our results (**Figure 3**). A direct visualization of the gene distribution revealed that the majority of genes formed one single cloud while several isolated groups of genes scattered around. We extracted these gene islands and found they were mainly non-protein-coding genes. Island 2 was significantly populated with the snoRNA genes (pink dots, p=1.07×10^−72^, Fisher’s Exact Test). Island 4, located to the very right of the plot, mainly contains human cDNA/PAC clone genes. microRNA genes (cyan dots) were mainly distributed in island 2 (p=3.99×10^−19^), island 4 (p=3.51×10^−73^), and island 5 (p=2.64×10^−41^). A group of ncRNAs which start with “LOC” and are often uncharacterized split the whole distribution into the left panel and the right panel (red dots, **Figure 3**). In the left panel, we observed a cluster of open reading frames (yellow dots, **Figure 3**) in the human genome.

**Figure 3:**
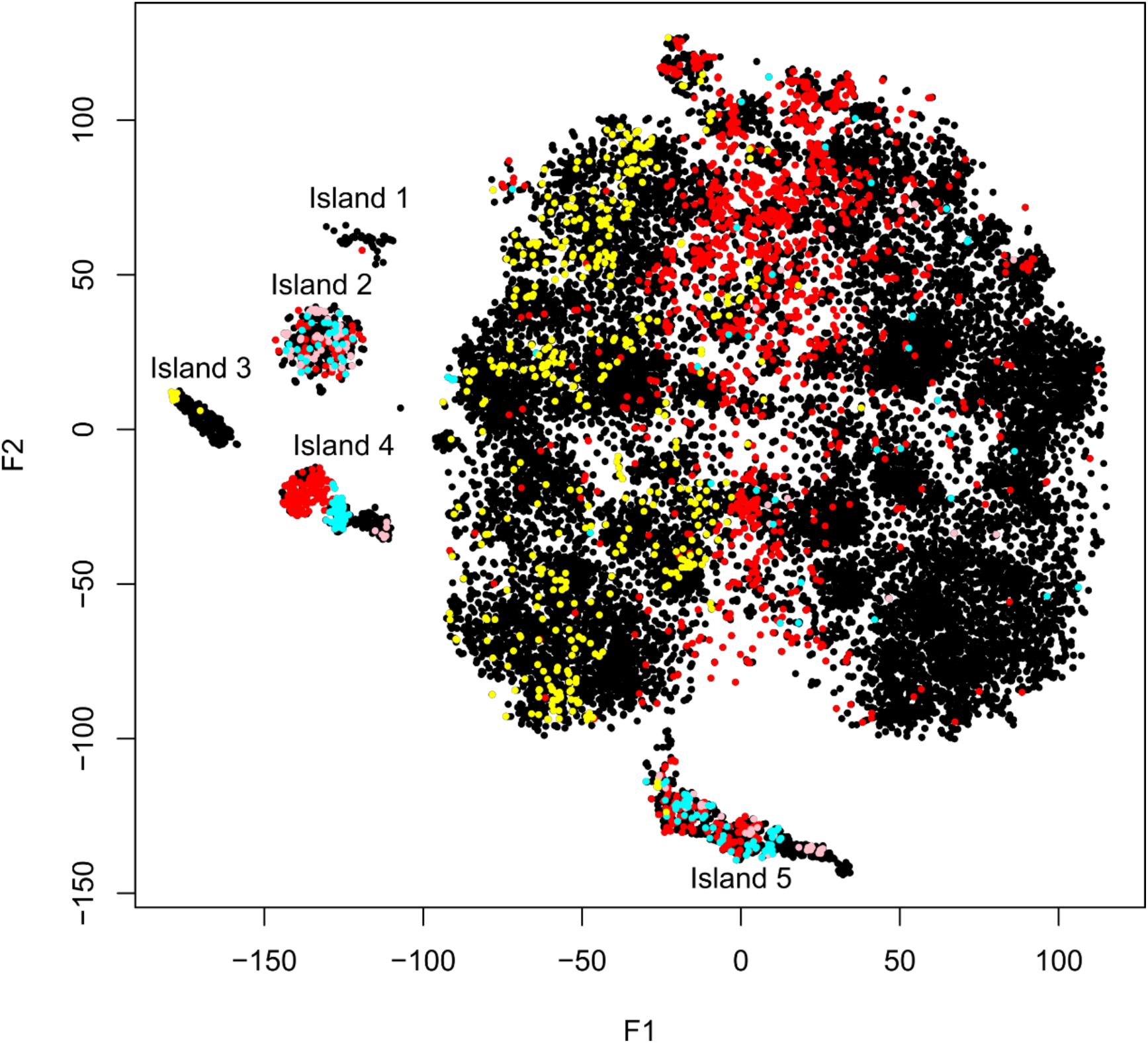
Gene co-expression map generated from embedding reveals clusters of functionally related genes. F1 and F2 are the first and the second dimensions of t-SNE. Red: LOC non-coding genes; cyan: microRNA; pink: small nucleolar RNA (snoRNA); yellow: undercharacterized ORFs.

### Tissue specific genes form spatial patterns in gene embedding

We mapped genes with z-scores representing their tissue-specific expression onto the gene co-expression map. We observed clear clusters in several tissues such as blood, skin, spleen, and lung (**Figure 4** and **supplementary figures**). Genes with high tissue specificity in blood highlighted two distant clusters. This is likely because that blood samples are relatively more widely used in gene expression studies and blood-specific genes and their relationships are thus better represented in our map. Tissues that are biologically relevant showed similar patterns. For example, tissues of female reproductive systems presented graded and similar patterns, including breast, ovary, and uterus. In these tissues, genes located in the bottom part of the map in general showed increased tissue specificity, compared to genes located on the top part of the map (**Figure 4** and **supplementary figures**). Interestingly, we found a group of ribosomal genes (~50) that were highly expressed in ovary and formed a small cluster in our map. In addition, cognition and neurology related tissues, such as brain (**Figure S5**), nerve (**Figure S14**), and pituitary (**Figure S17**), presented quite diverse patterns. Nerve and pituitary are more similar to each other, with a wide range of genes showing moderate tissue-specificity distributed across the whole map. In contrast, active genes in brain, which are mainly distributed on the top part of the map, are much smaller in numbers but showed much stronger tissue-specificity (red dots, **Figure S5**). Notably, all tissues except the blood are expected to be under-represented in the GEO data we used because tissue samples are difficult to obtain for human.

**Figure 4:**
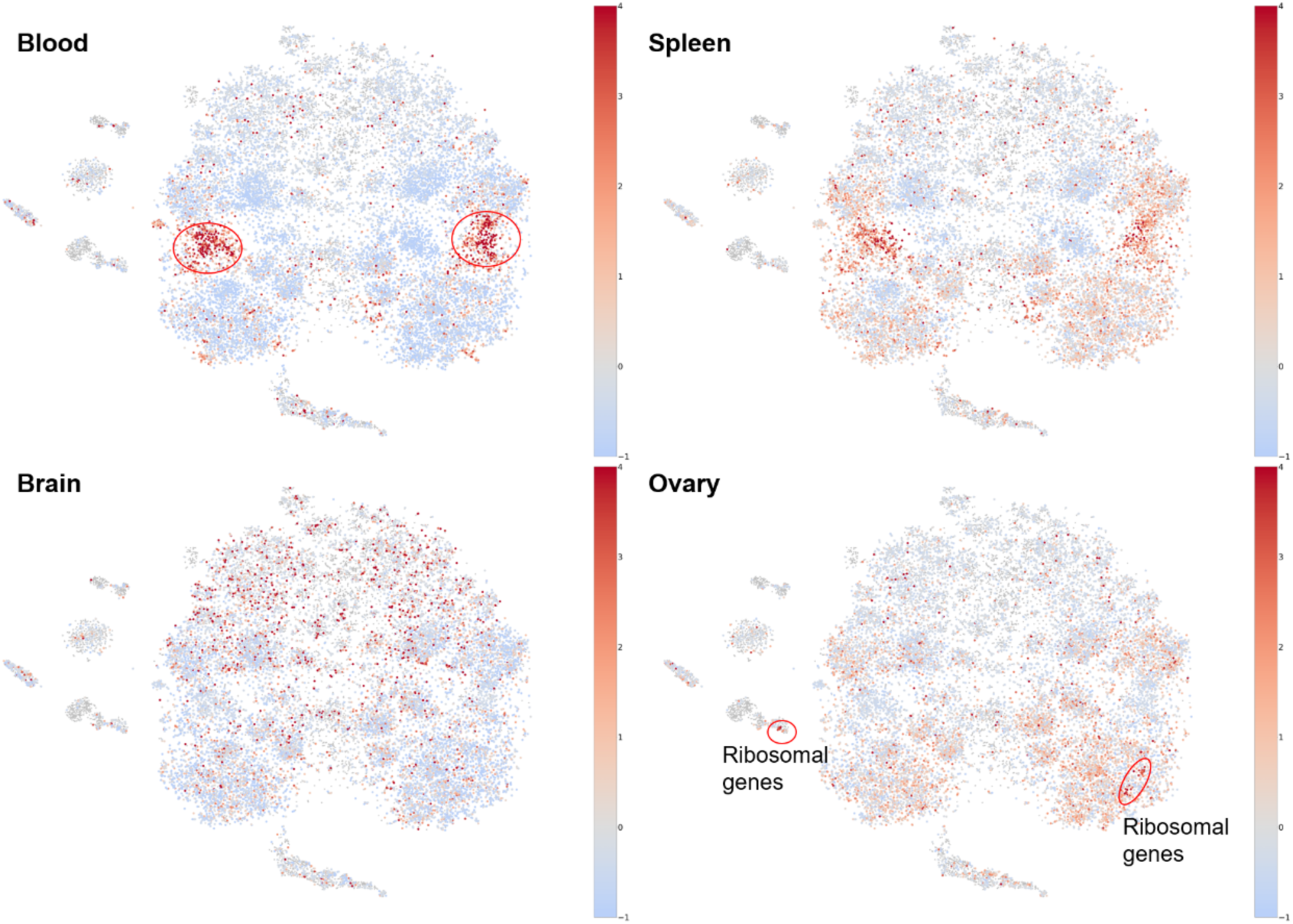
Embedding reveals clusters of genes with tissue-specificity. Blood and spleen have clear patterns of tissue-specific genes. Reproductive system (e.g., ovary) also showed distinguished genes. Genes not available in GTEx data were colored grey.

### Prediction of gene-gene interaction using embedded vectors

The performances of GGIPNN with embedding matrix are presented in **Figure 5**. Using gene embedding matrix derived from GEO but do not make them trainable, we achieved an AUC of 0.720 over the test set, in which there are no gene overlapping with the training set nor the validation set. The AUC score is lower, 0.664, for the GGIPNN with gene embedding matrix derived from GEO as initial weights but trainable. This is understandable as the gene embedding matrix for the genes in the training set was updated and leaving the gene embedding matrix in the test set “out of sync” with that for the training set, i.e., overfitting. As expected, the GGIPNN with both untrainable and trainable random embedding matrix have AUC scores (0.505 and 0.493) close to random (0.5).

**Figure 5:**
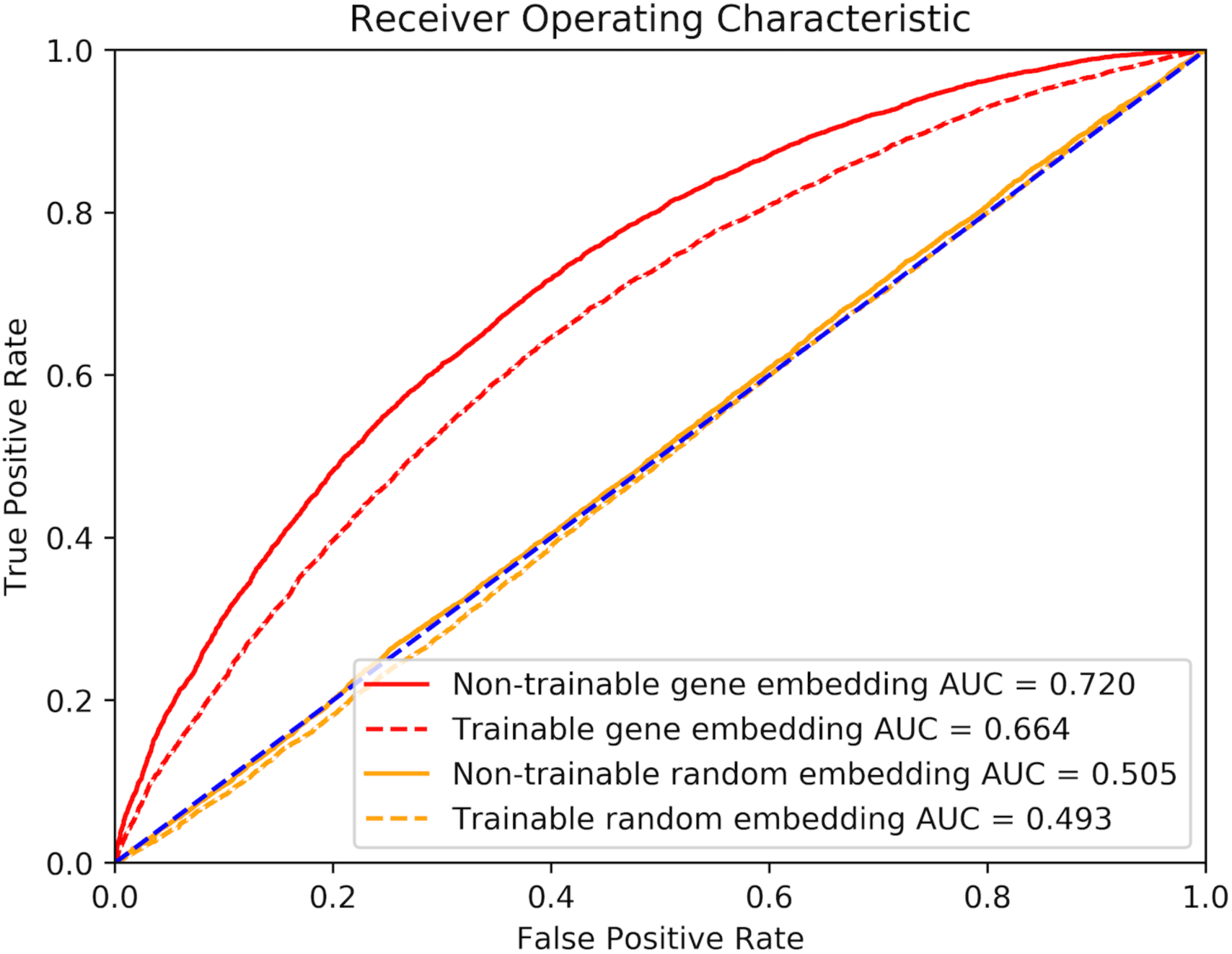
ROC curves for gene-gene interaction predictor neural nestworks.

## DISCUSSION

In this work, we explored the idea of distributed representation of genes using their co-expression. Purely trained from their co-expression patterns in GEO, except using MSigDB as hyper-parameter tuning, the trained embedding matrix captures functional relationships among genes. In the t-SNE generated gene co-expression map of the embedding matrix, tight clusters of non-coding genes are formed, while broader clusters corresponding to tissue specific genes are also visible.

The usefulness of gene embedding is beyond simply a nice visualization. Using the gene embedding as the basic layer for a multi-layer neural network, we can predict the gene-gene interaction with an AUC of 0.720. This is an intriguing result because the only input to the predictor is the names of the two genes. Therefore, the distributed representation of the genes, i.e., their embeddings, are laden with rich semantic information about their function.

The concept of concept embedding is not new to molecular biology. Works had been done to geometrical embedding gene co-expression networks into 2-D planar networks [18]. Recently, in the spirit of embedding everything, the work of bioVectors have been developed to embedding kmers in biological sequences into distributed representation [19]. Yang et al leveraged the Doc2vec model to learn embedded representations of protein sequences [20]. Similarly, a project named ‘Gene2vec’ is available embedding gene sequences [21]. However, to the best of our knowledge, our work is the first to directly embed genes into distributed representations based on their natural context - their expression and co-expression.

In this work, we are using the gene co-expression as the definition of “context” for gene embedding. However, it is possible to extend the current work to include other definitions of context for genes. For example, co-occurrence of genes across species, gene-gene and protein-protein interactions from experiments, and co-occurrences of genes in literature, all can be a source of information to define context.

The distributed representation of genes can enable new applications. E.g., as illustrated in the Results, with continuous representation of genes, it is possible to direct feed gene as inputs to neural networks, and can be useful for any prediction tasks with gene names as input.

A limitation of current approach is the lack of higher order semantics. In word embedding a surprising result was that the direction of the embedding space can be interpreted. For example, the vector representations of the words King, Queen, Man, and Woman formed a parallelogon. This higher order of semantics from NLP modeling may be due to that the concepts between these words were connected by certain relationships, which is reflected by the occurrences of these words being connected by certain verbs. To achieve this level of semantic embedding of genes, future works modeling more information about genes are warranted.

## CONCLUSIONS

We proposed a machine learning method that utilizes transcriptome-wide gene co-expression to generate a distributed representation of genes. We further demonstrated the utility of our distribution by predicting gene-gene interaction based solely on gene names. We believe that this distributed representation of genes could be useful for more bioinformatics applications.

## Declarations

### Ethics approval and consent to participate

N/A.

### Consent for publication

Not applicable

### Availability of data and material

The codes for gene2vec training, visualization, and gene-gene interaction prediction are available on GitHub (https://github.com/jingcheng-du/Gene2vec). We also released our pre-trained gene2vec on GitHub (https://github.com/jingcheng-du/Gene2vec/blob/master/pre_trained_emb/gene2vec_dim_200_iter_9.txt).

### Competing interests

The authors declare that there is no competing interest.

### Funding

Research was partially supported by the Cancer Prevention Research Institute of Texas (CPRIT) Training Grant #RP160015.

### Disclaimer

The content is solely the responsibility of the authors and does not necessarily represent the official views of the Cancer Prevention and Research Institute of Texas.

### Authors’ contributions

JD, PJ and DZ designed the study, performed the experiments and drafted the manuscript. YD, and CT assisted to the study design. ZZ and DZ supervised the study. Everyone read and revised the manuscript.

## Acknowledgments

We thank the anonymous reviewers for their careful reading of our manuscript and their many insightful comments.

