## Supplementary Materials for "Gene2Vec: Distributed Representation of Genes Based on Co-Expression" (from another version of this preprint: version 1, 2018-03-22)

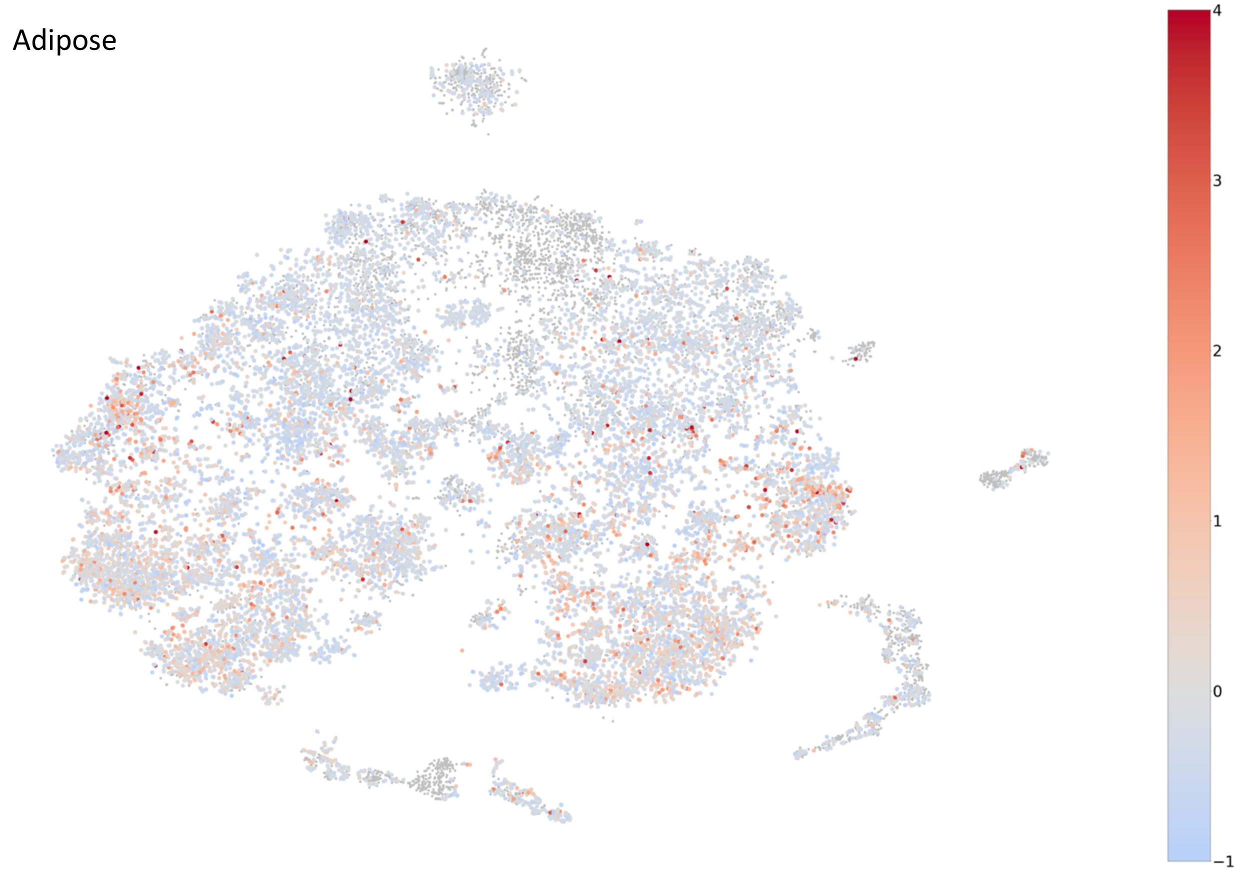


**Figure S1**: Gene co-expression map with genes colored by z-scores obtained using GTEx adipose expression data.

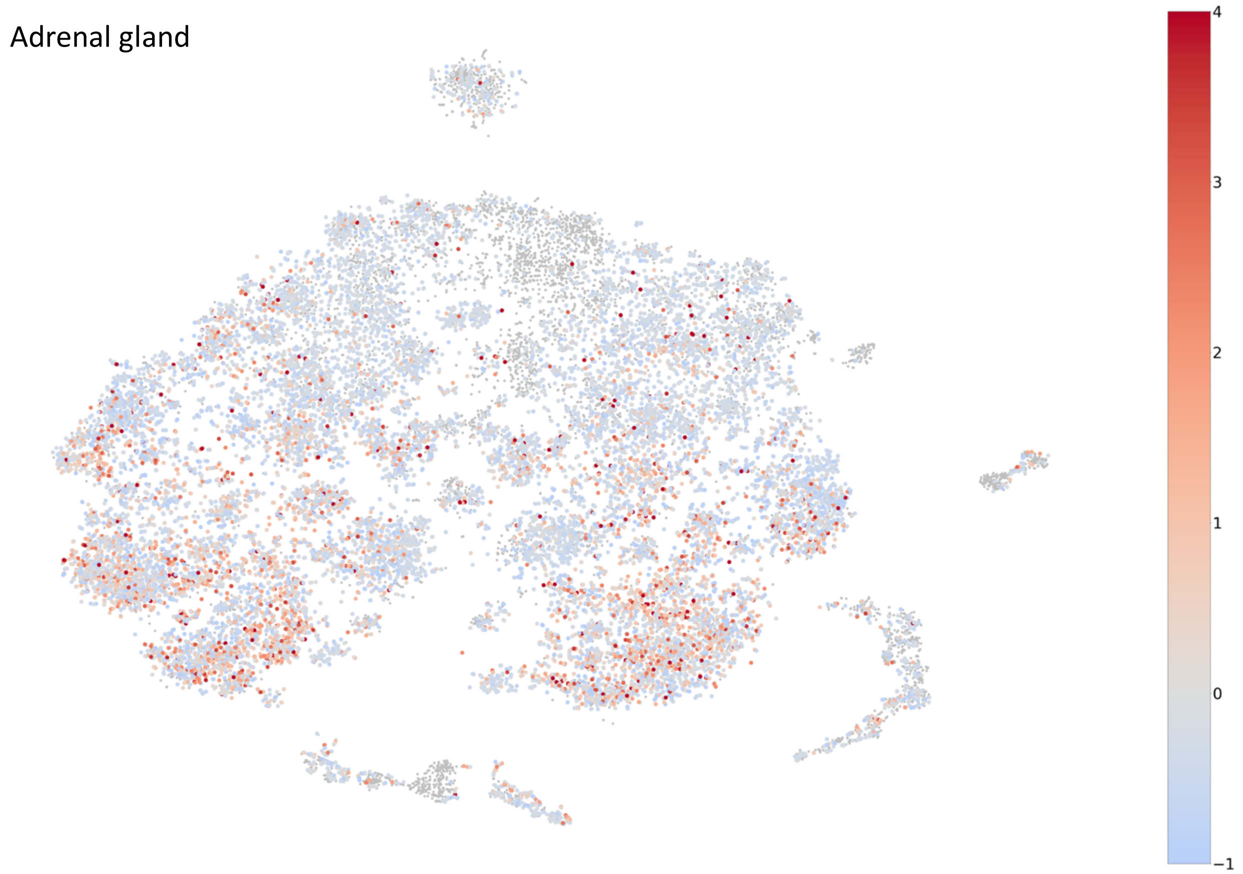


**Figure S2**: Gene co-expression map with genes colored by z-scores obtained using GTEx adrenal gland expression data.

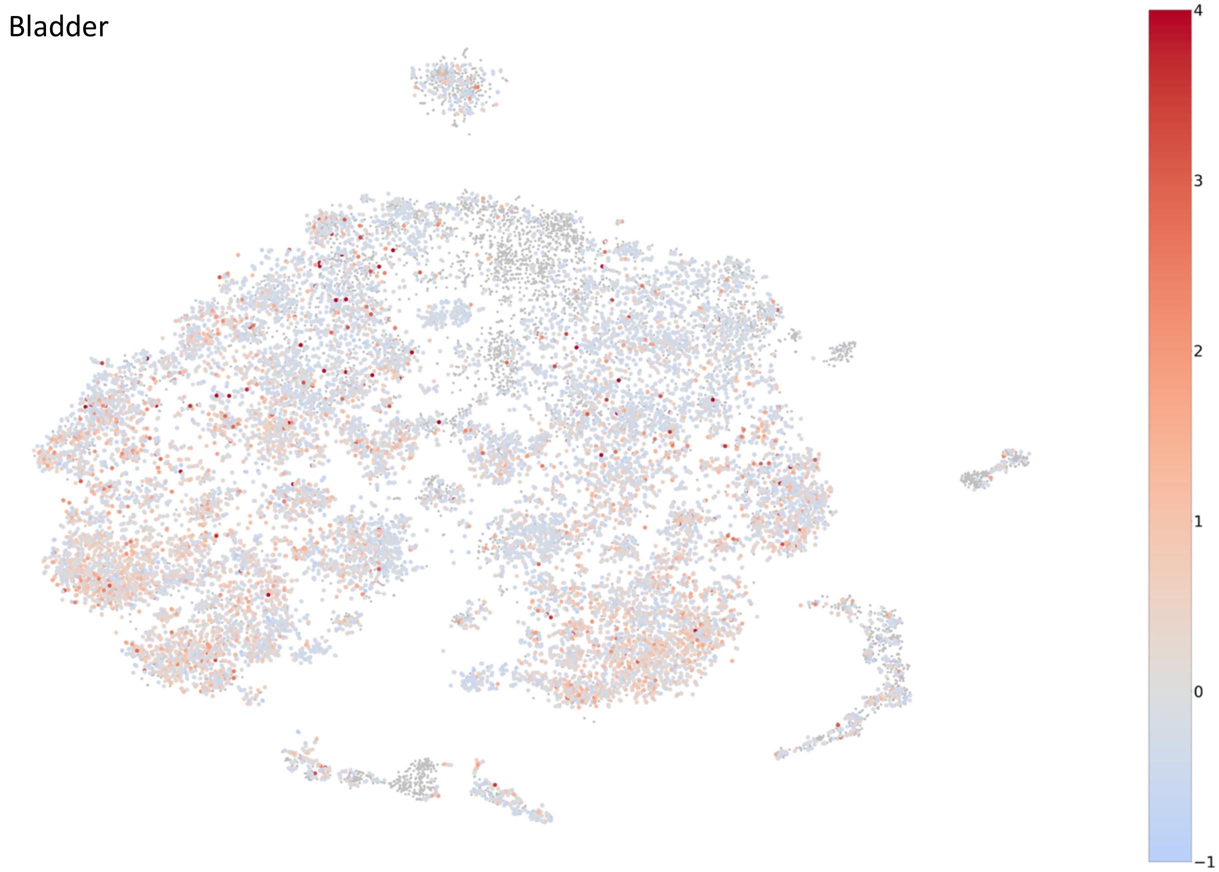


**Figure S3**: Gene co-expression map with genes colored by z-scores obtained using GTEx bladder expression data.

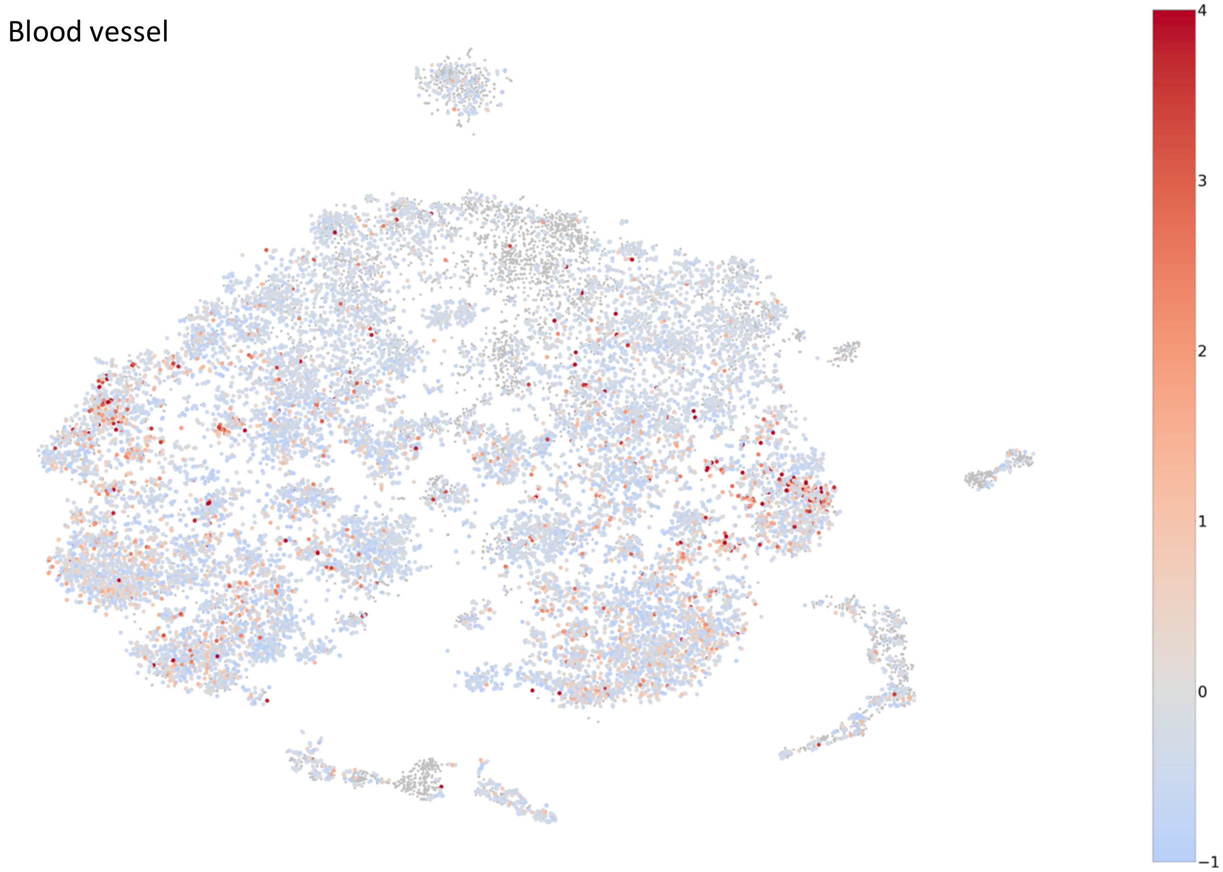


**Figure S4**: Gene co-expression map with genes colored by z-scores obtained using GTEx blood vessel expression data.

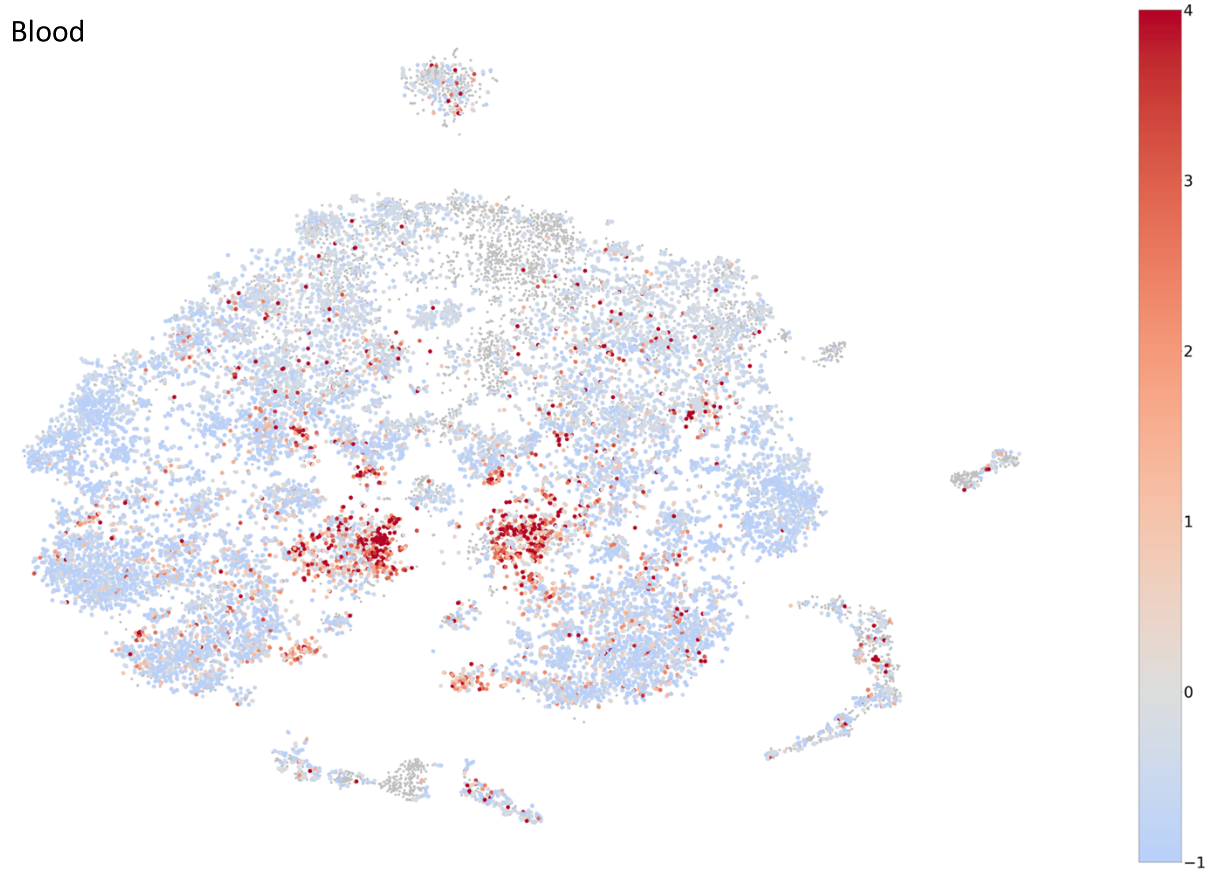


**Figure S5**: Gene co-expression map with genes colored by z-scores obtained using GTEx blood expression data.

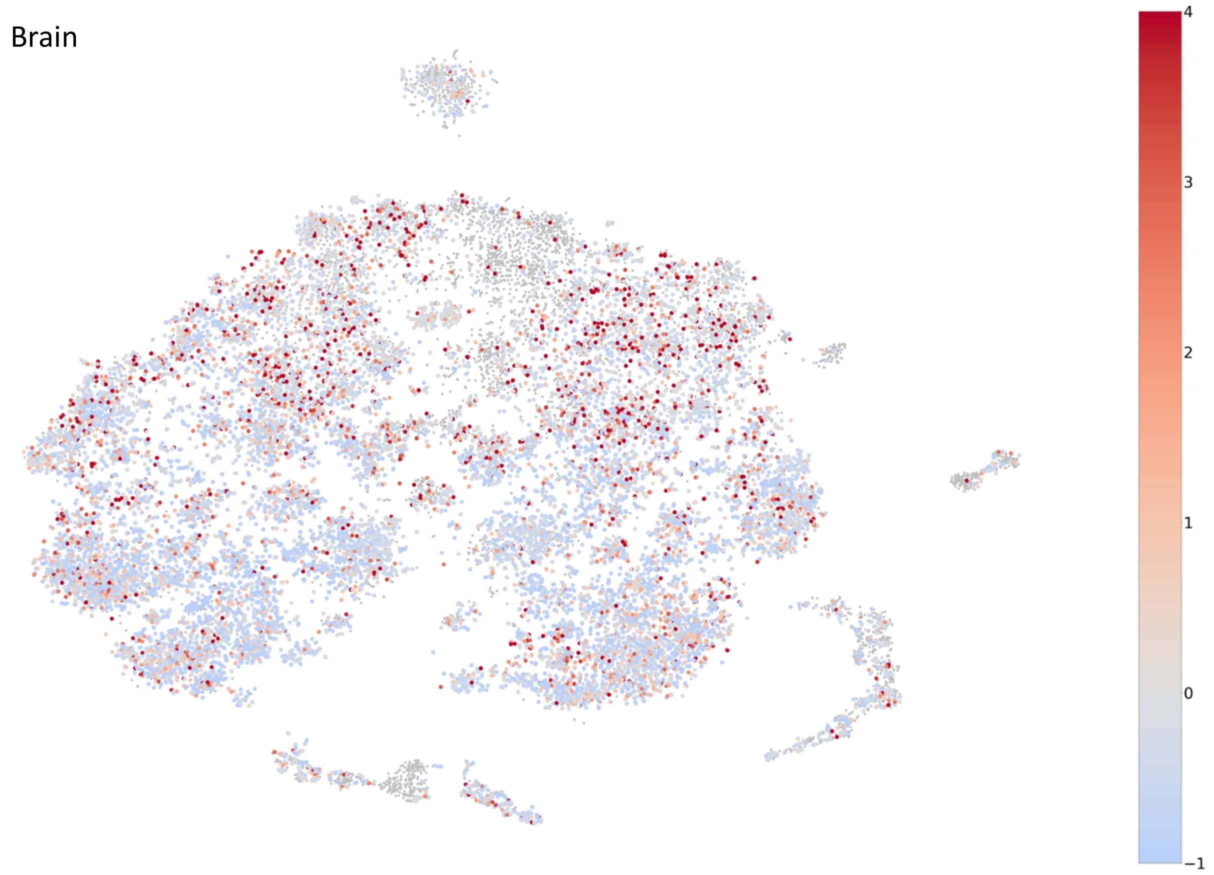


**Figure S6**: Gene co-expression map with genes colored by z-scores obtained using GTEx brain expression data.

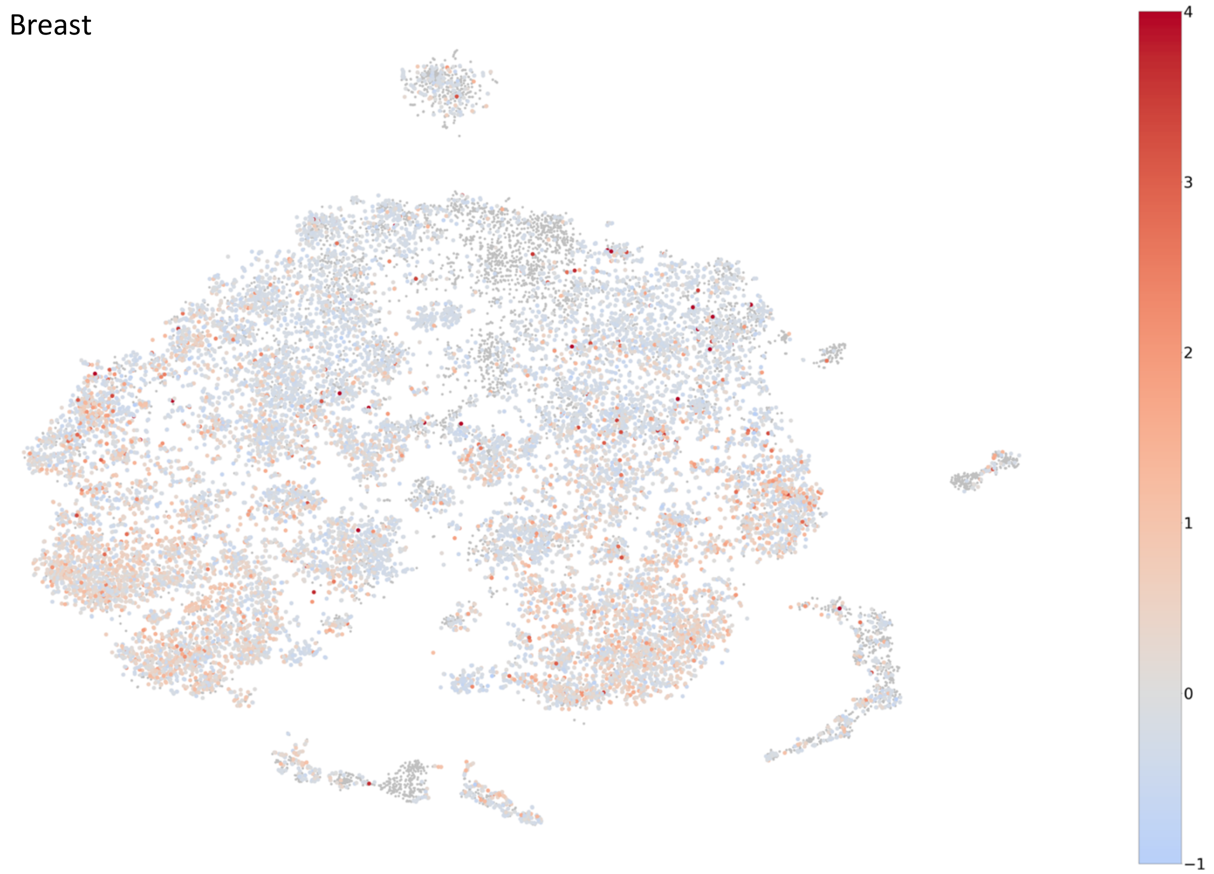


**Figure S7**: Gene co-expression map with genes colored by z-scores obtained using GTEx breast expression data.

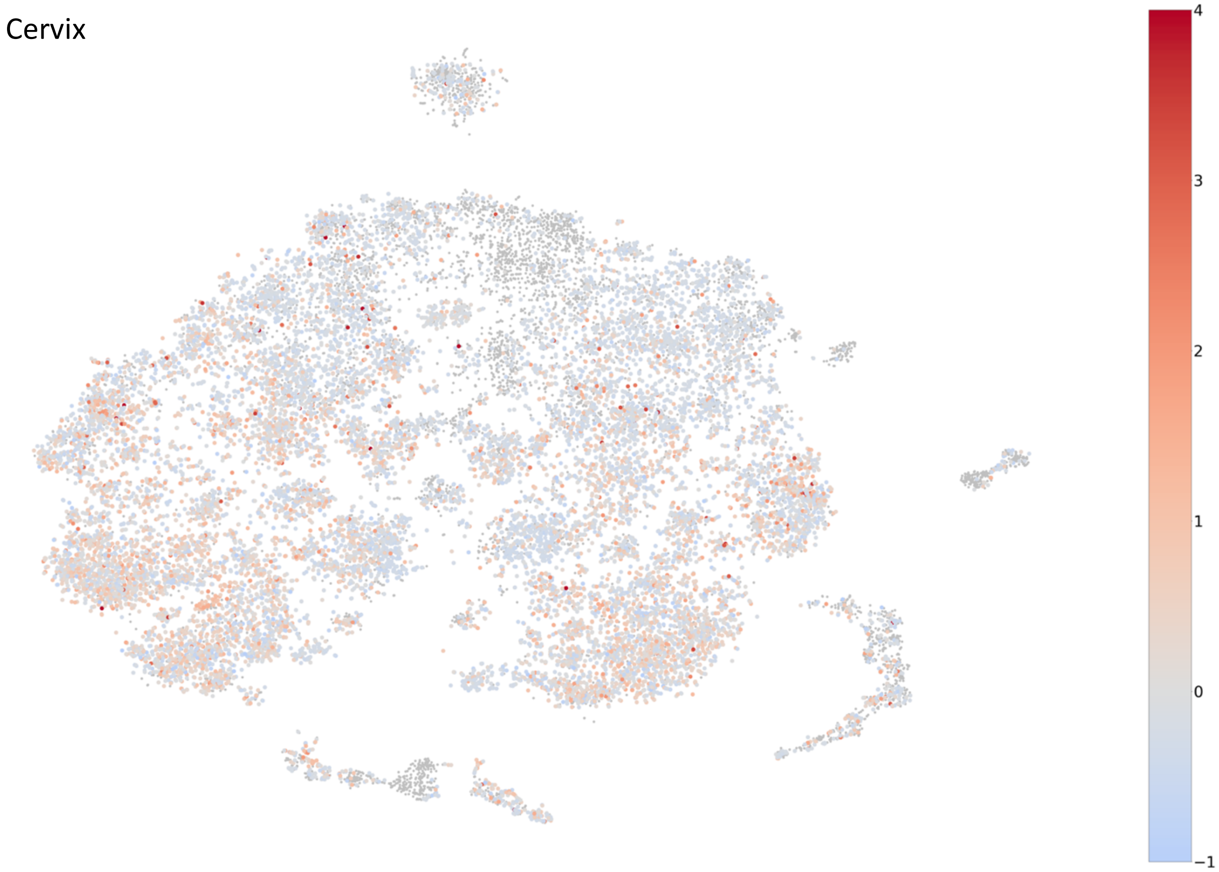


**Figure S8**: Gene co-expression map with genes colored by z-scores obtained using GTEx cervix uteri expression data.

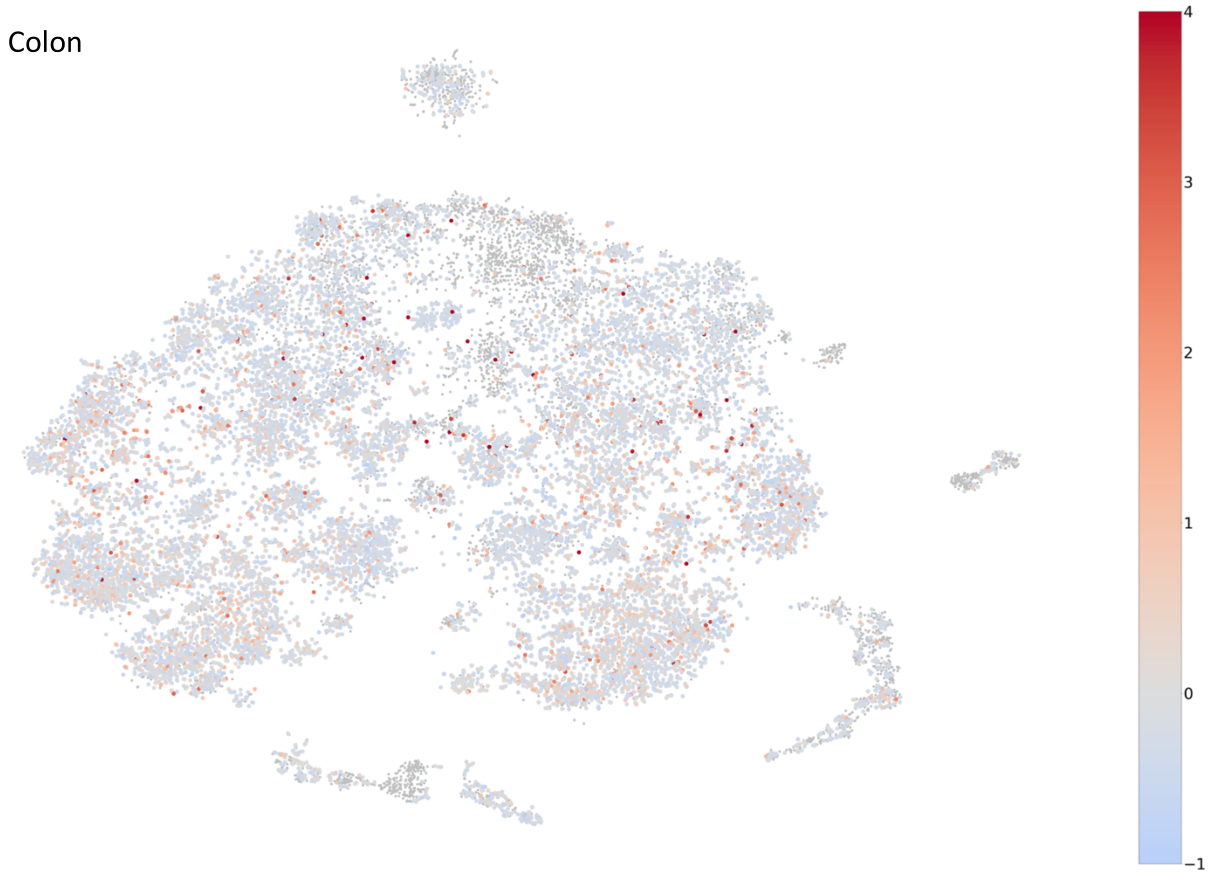


**Figure S9**: Gene co-expression map with genes colored by z-scores obtained using GTEx colon expression data.

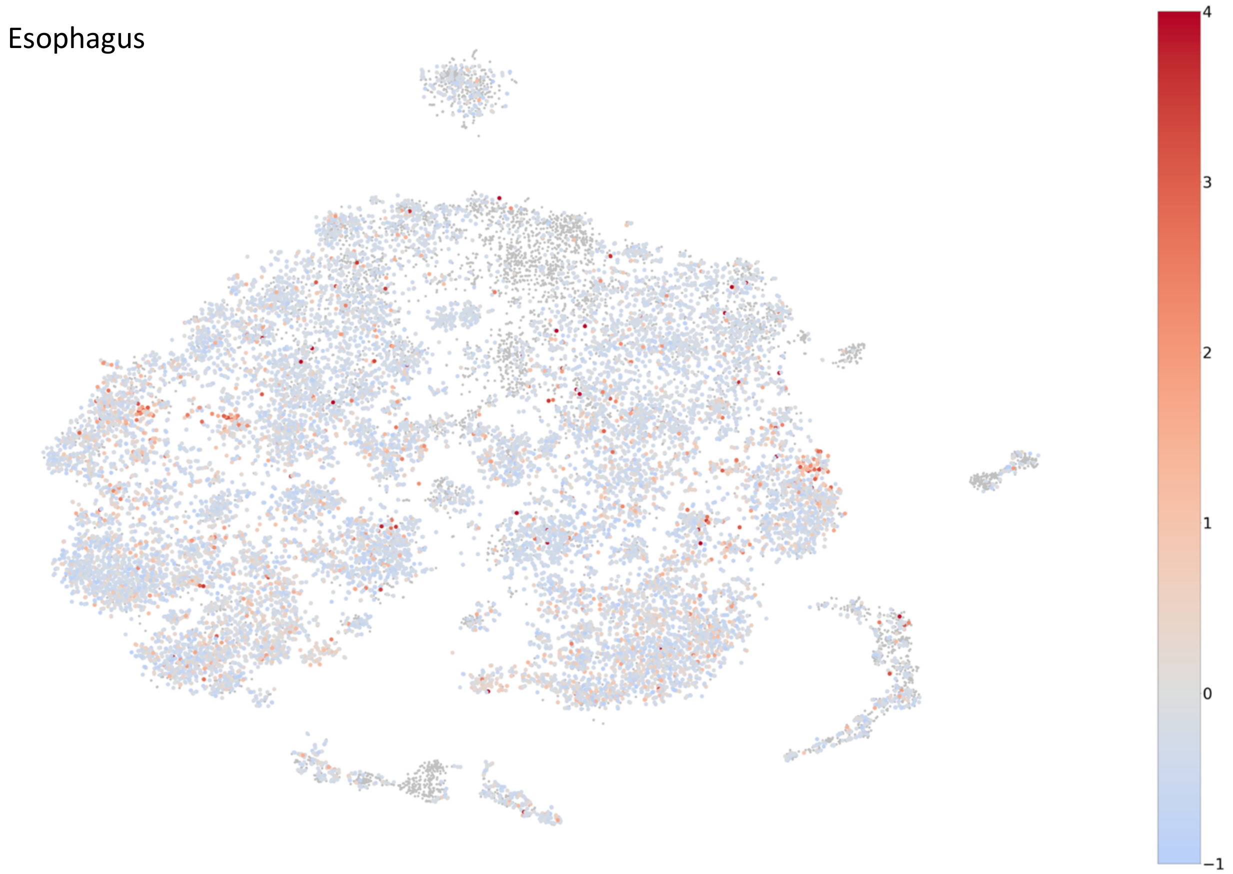


**Figure S10**: Gene co-expression map with genes colored by z-scores obtained using GTEx esophagus expression data.

Fallopian tube

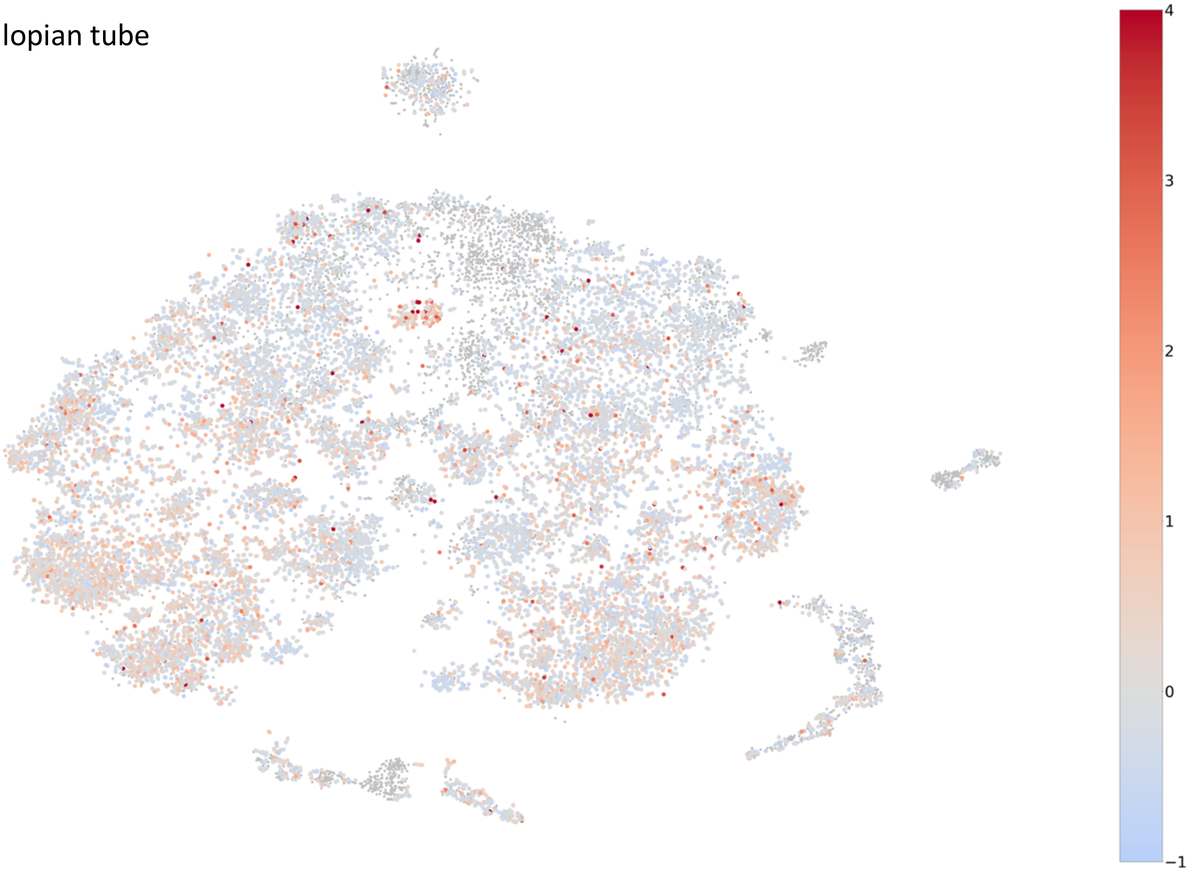


**Figure S11**: Gene co-expression map with genes colored by z-scores obtained using GTEx fallopian tube expression data.

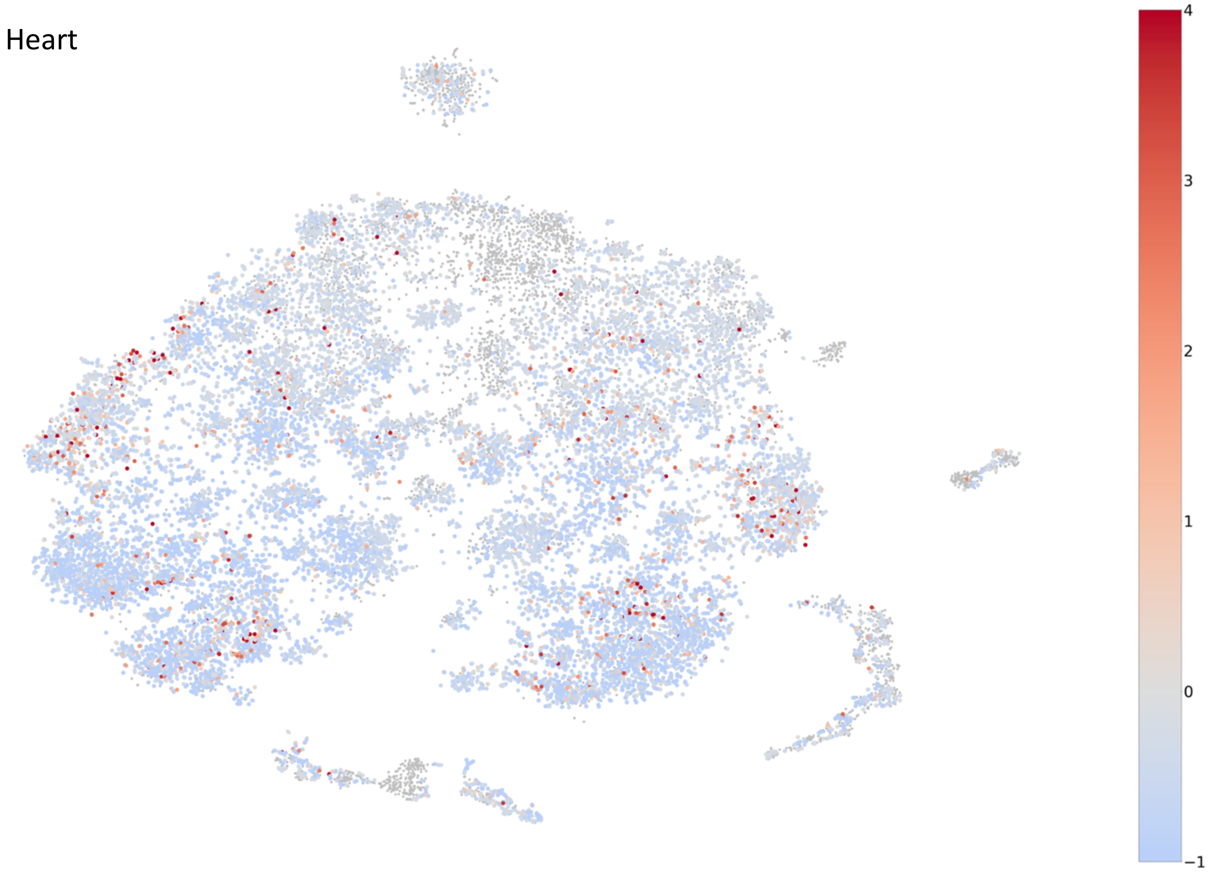


**Figure S12**: Gene co-expression map with genes colored by z-scores obtained using GTEx heart expression data.

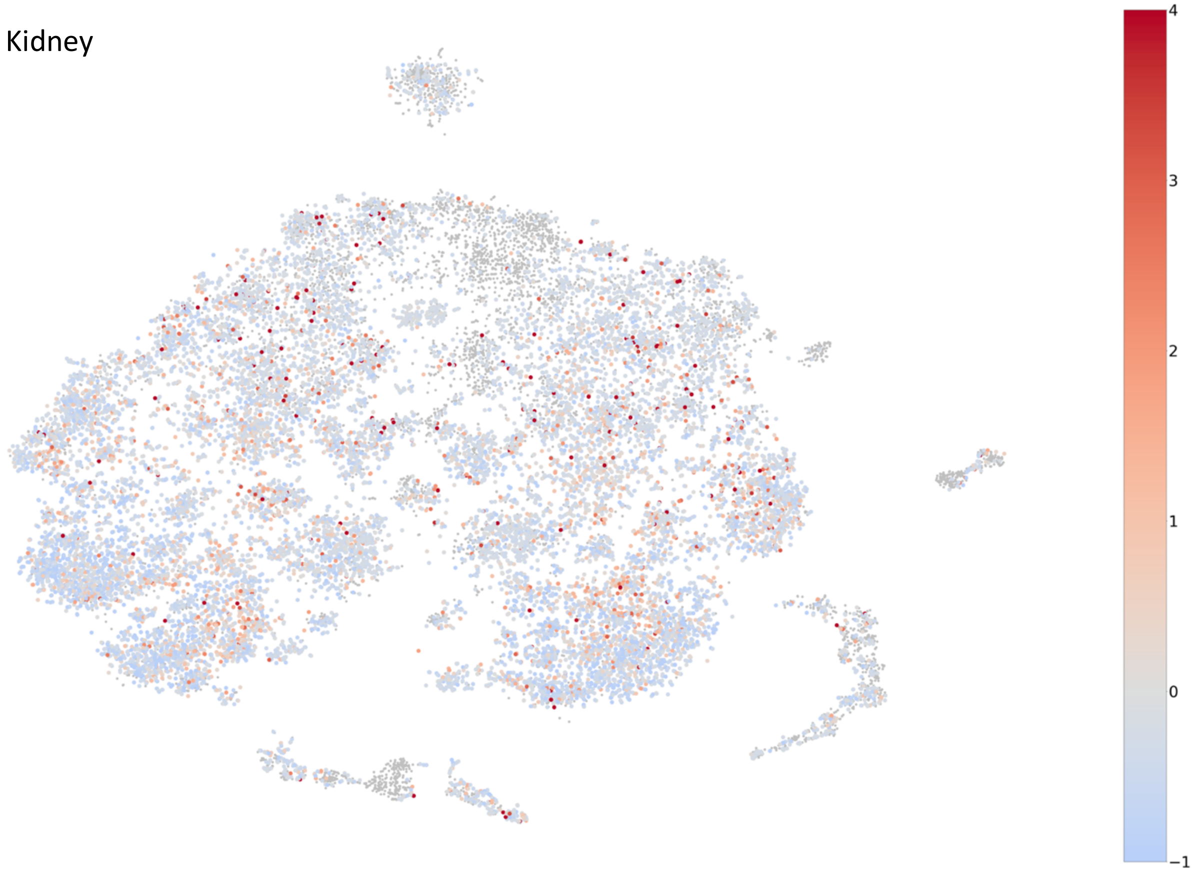


**Figure S13**: Gene co-expression map with genes colored by z-scores obtained using GTEx kidney expression data.

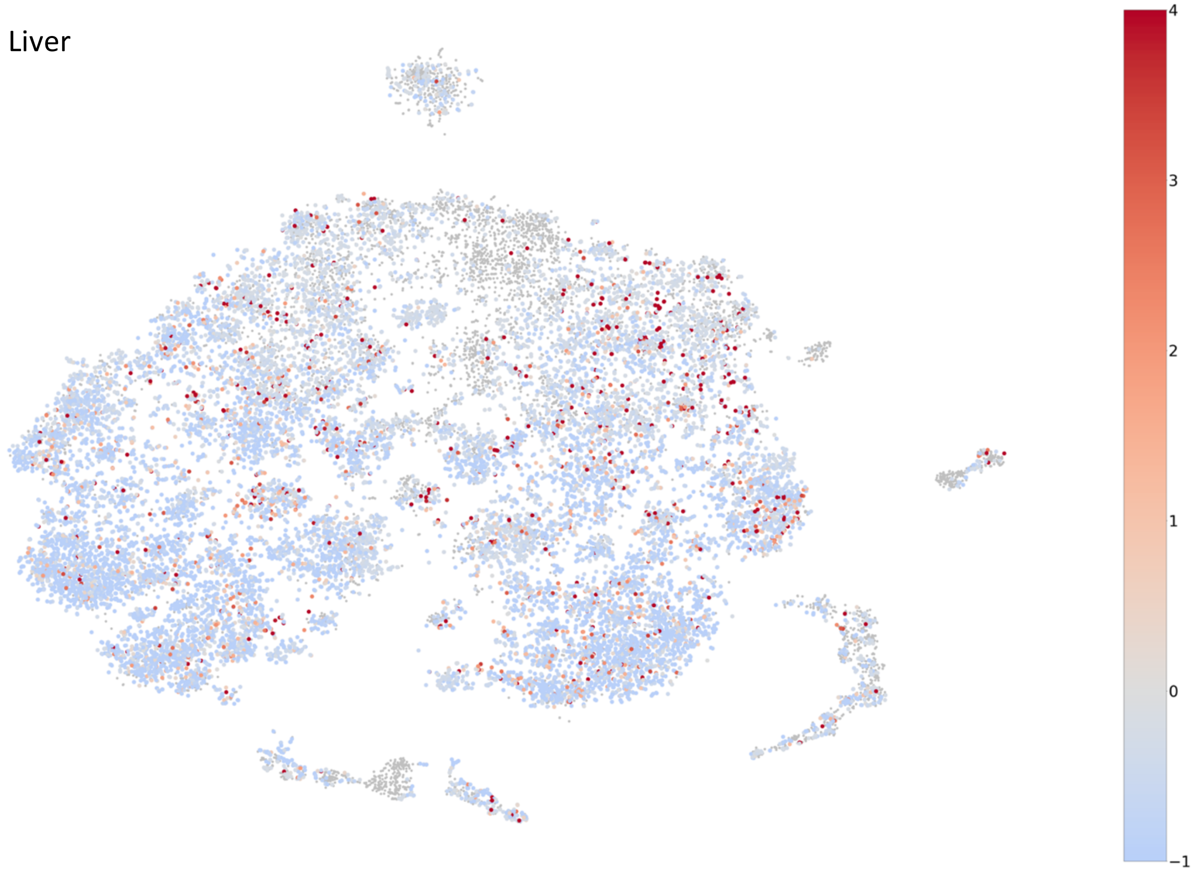


**Figure S14**: Gene co-expression map with genes colored by z-scores obtained using GTEx liver expression data.

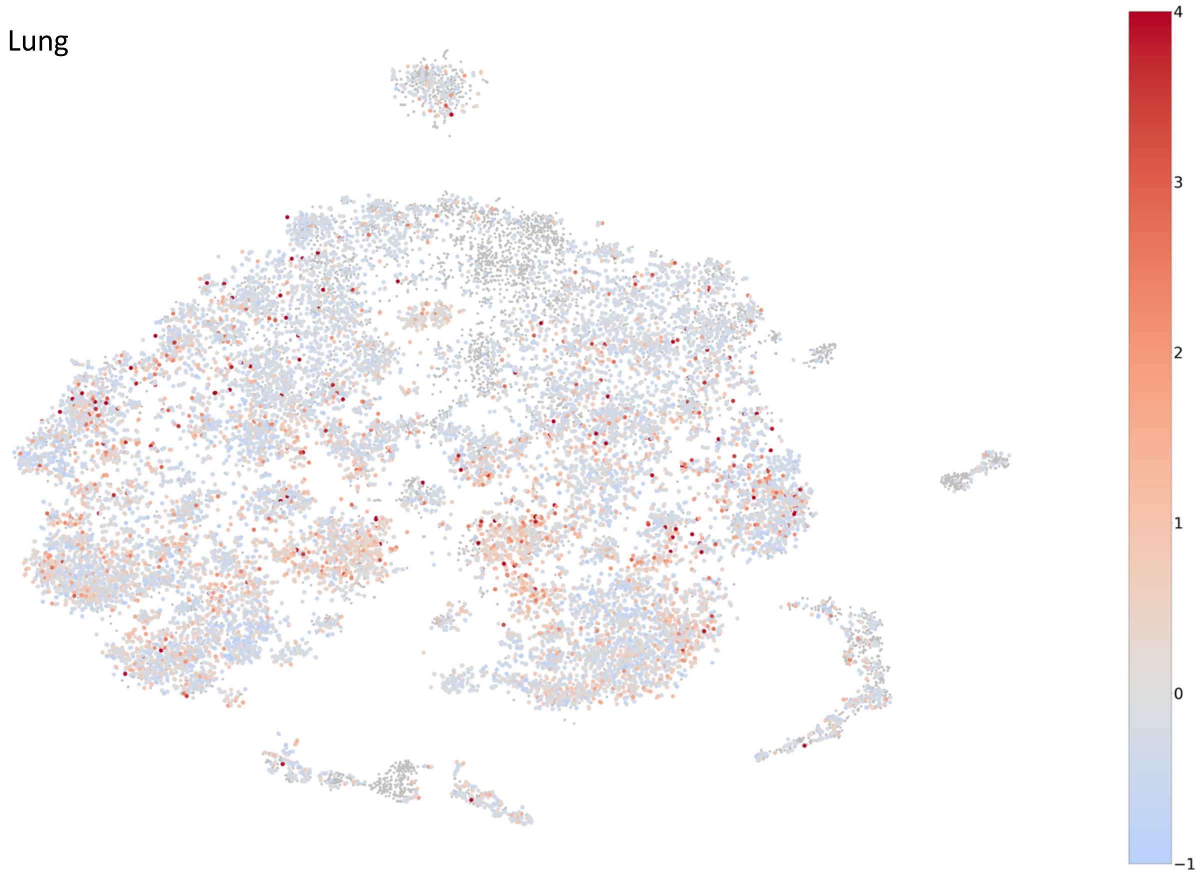


**Figure S15**: Gene co-expression map with genes colored by z-scores obtained using GTEx lung expression data.

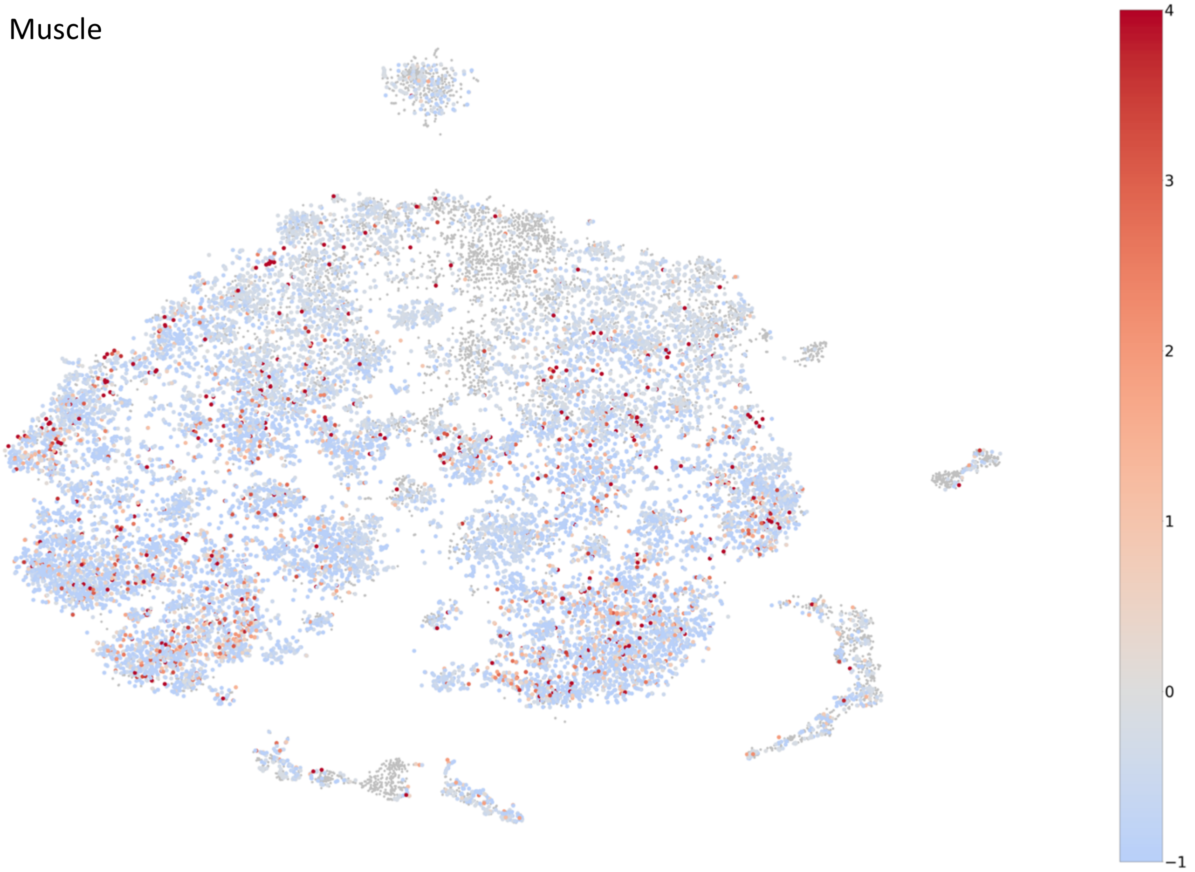


**Figure S16**: Gene co-expression map with genes colored by z-scores obtained using GTEx muscle expression data.

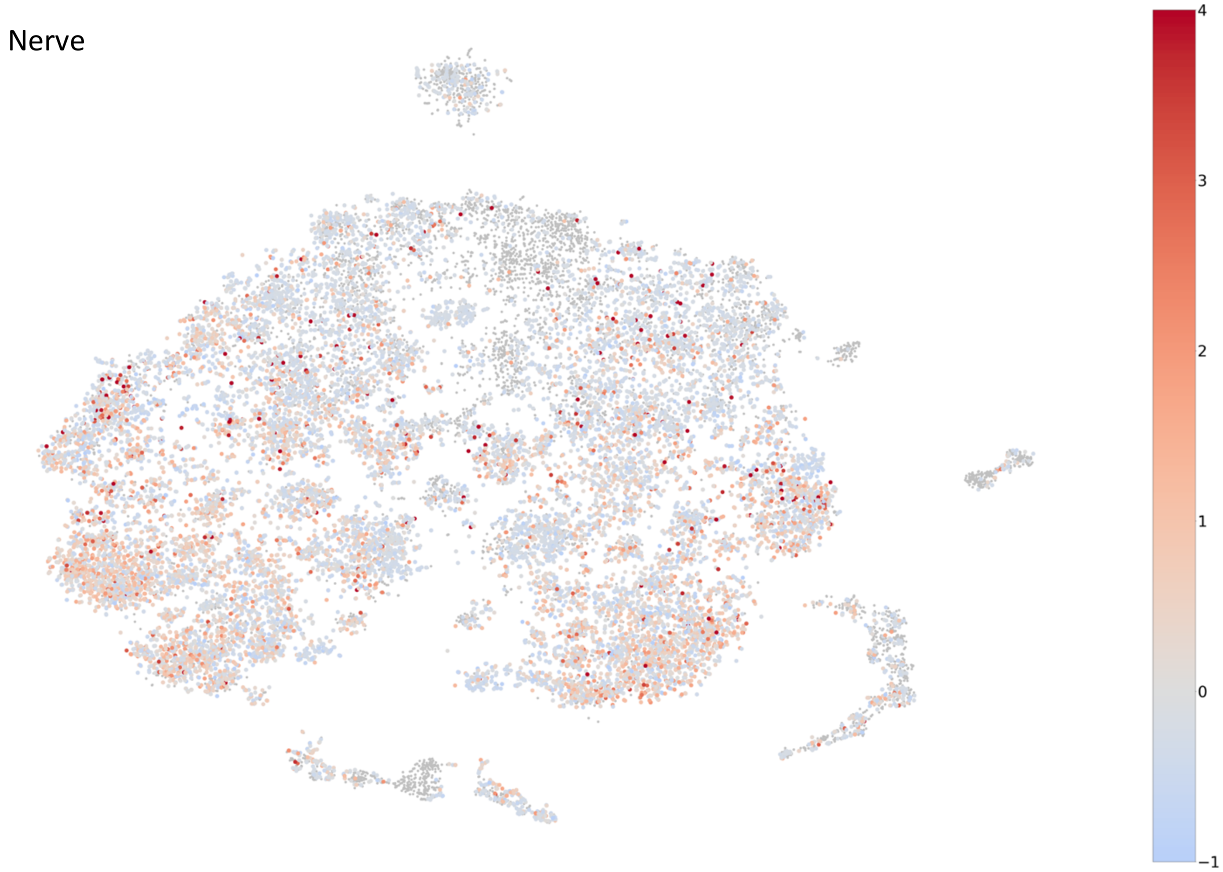


**Figure S17**: Gene co-expression map with genes colored by z-scores obtained using GTEx nerve expression data.

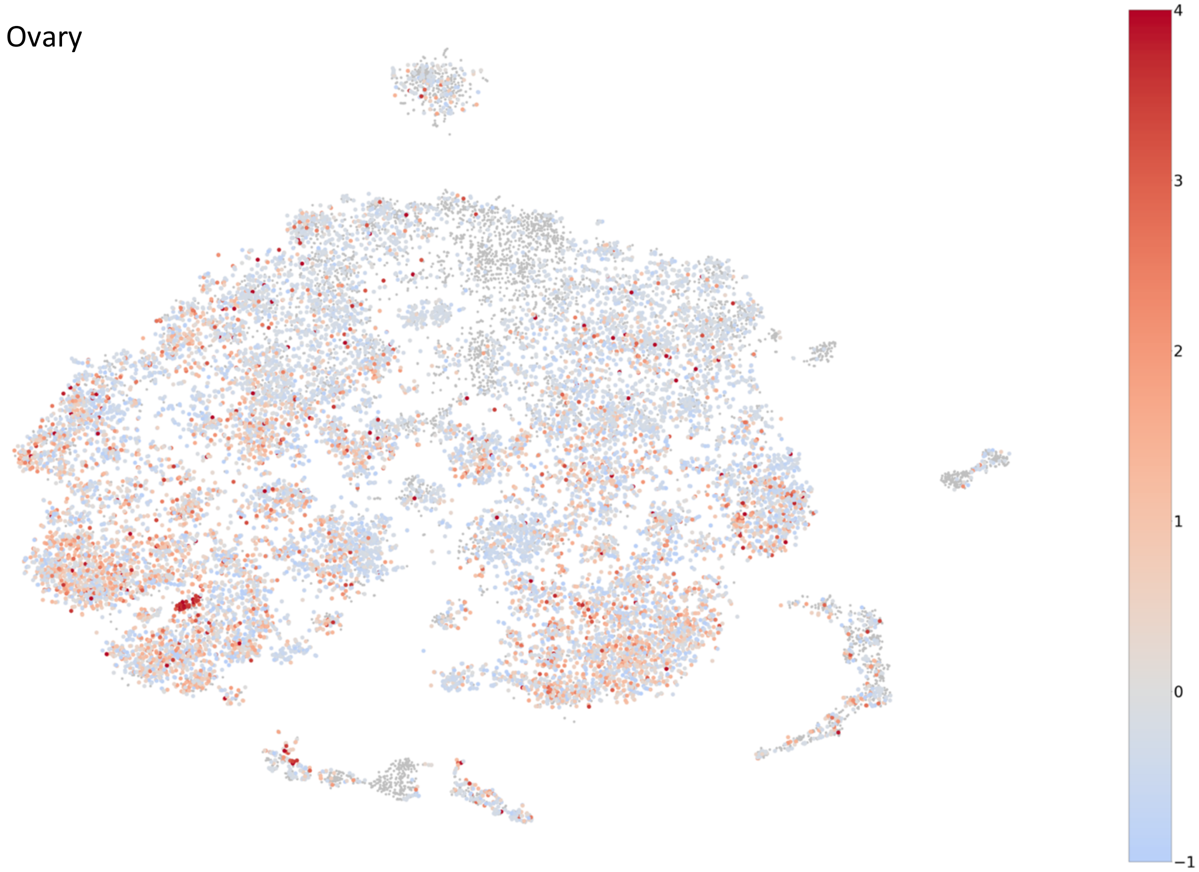


**Figure S18**: Gene co-expression map with genes colored by z-scores obtained using GTEx ovary expression data.

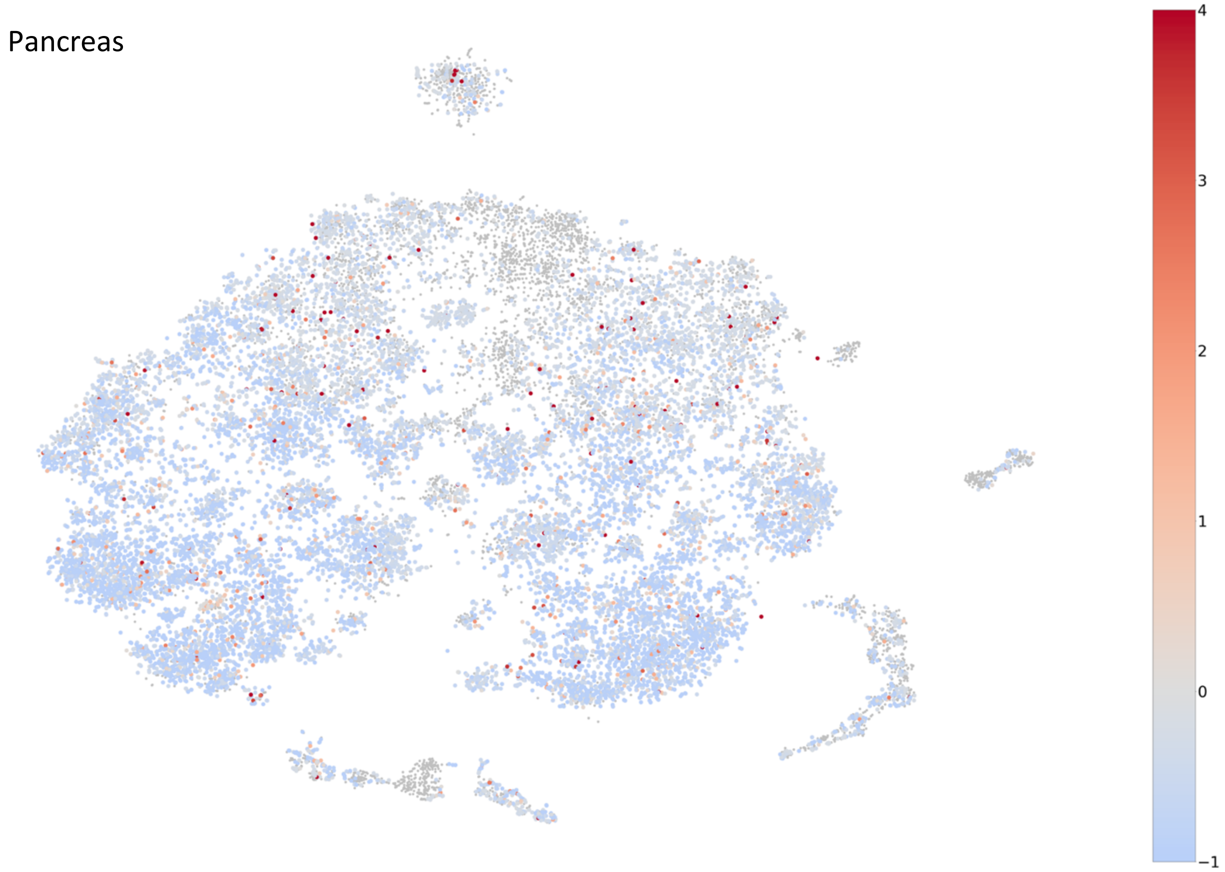


**Figure S19**: Gene co-expression map with genes colored by z-scores obtained using GTEx pancreas expression data.

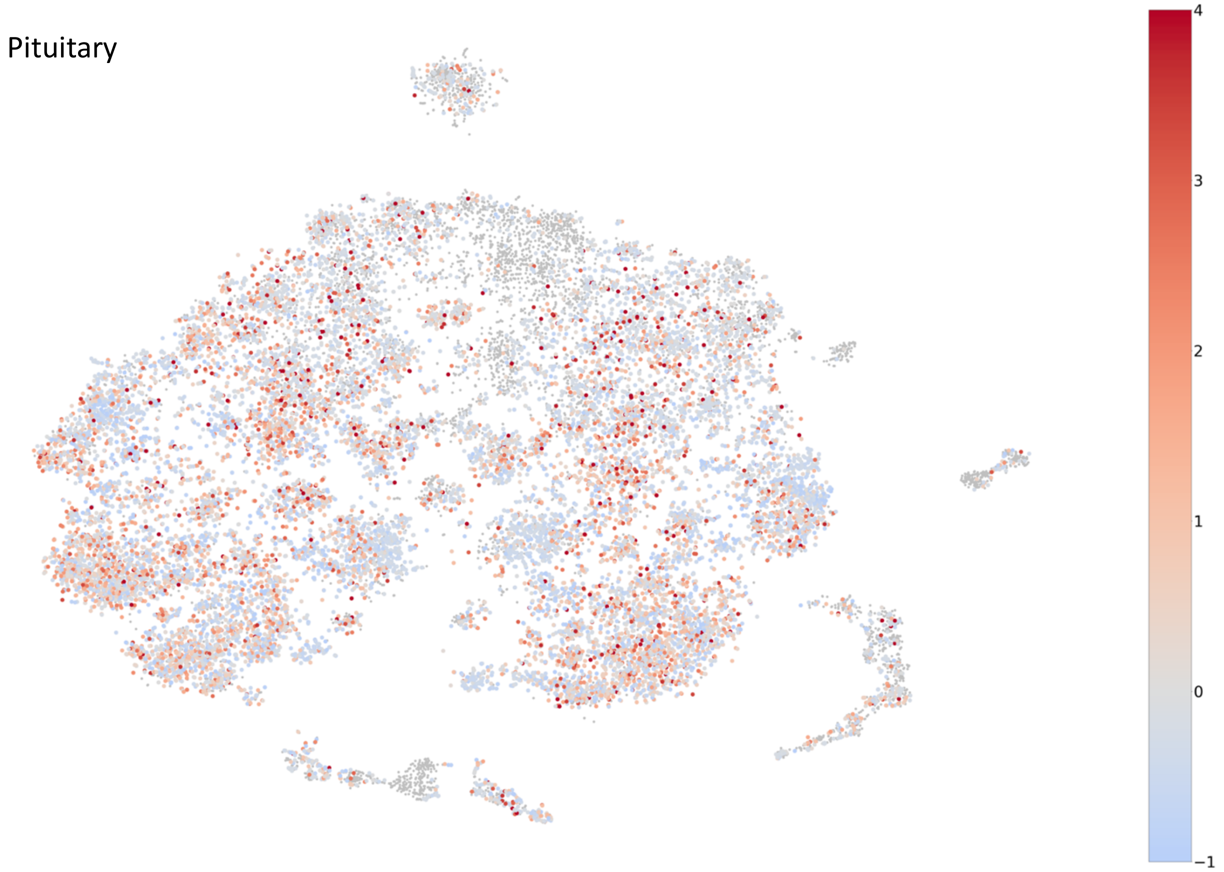


**Figure S20**: Gene co-expression map with genes colored by z-scores obtained using GTEx pituitary expression data.

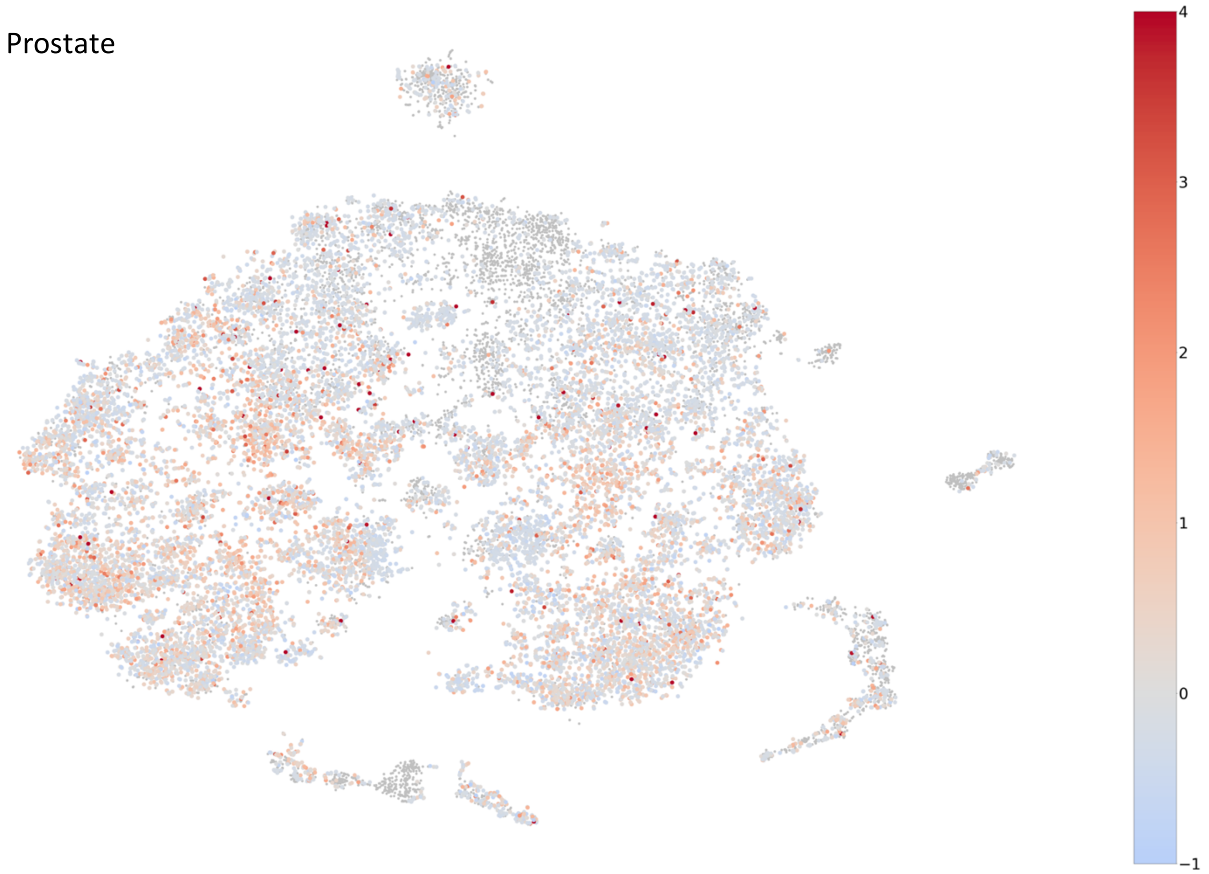


**Figure S21**: Gene co-expression map with genes colored by z-scores obtained using GTEx prostate expression data.

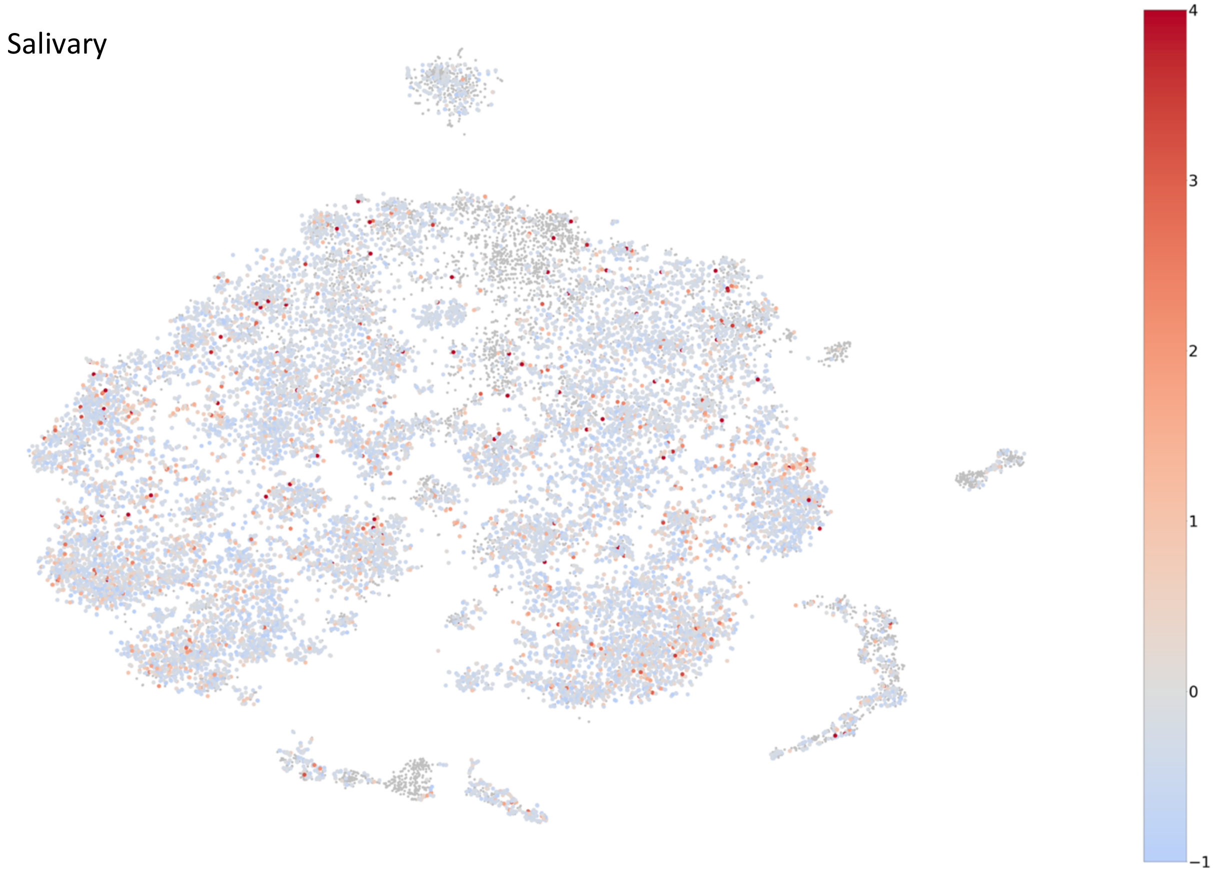


**Figure S22**: Gene co-expression map with genes colored by z-scores obtained using GTEx salivary gland expression data.

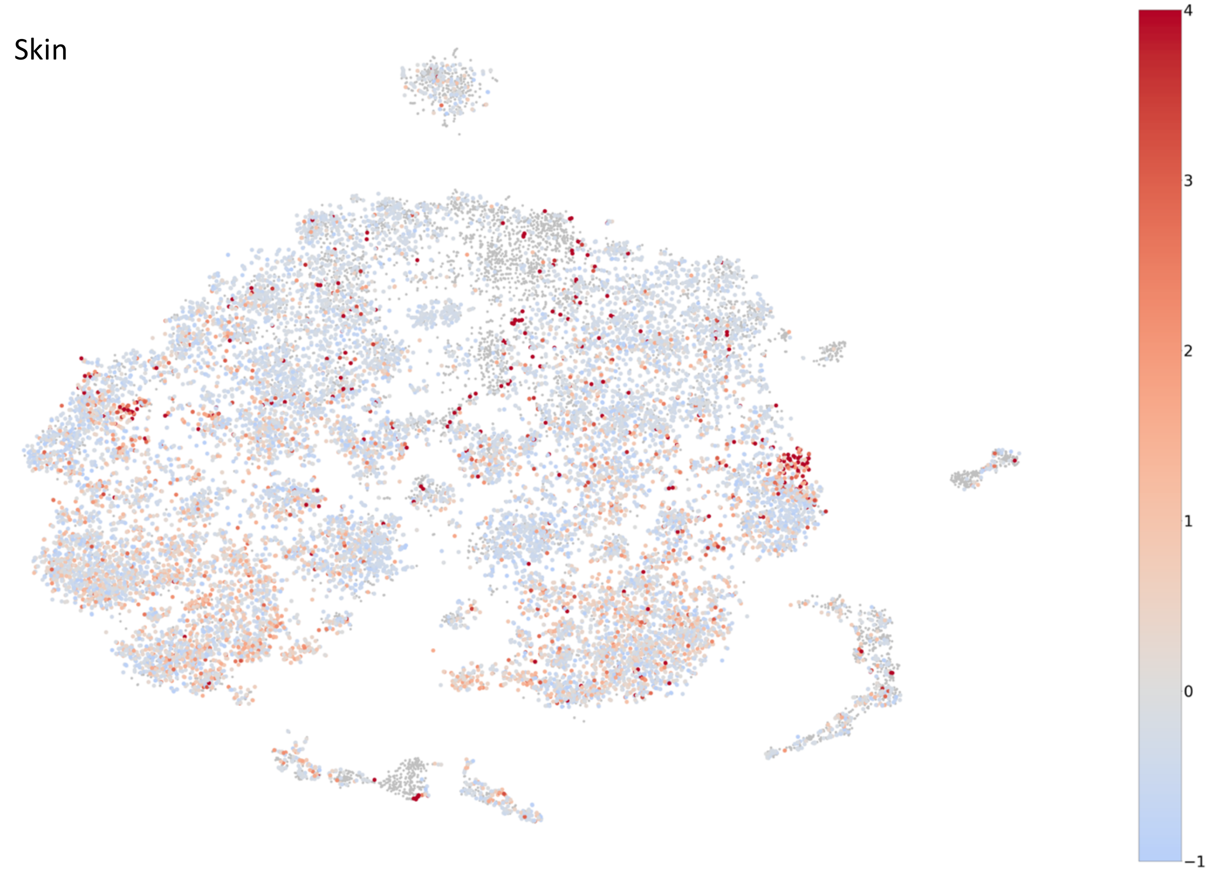


**Figure S23**: Gene co-expression map with genes colored by z-scores obtained using GTEx skin expression data.

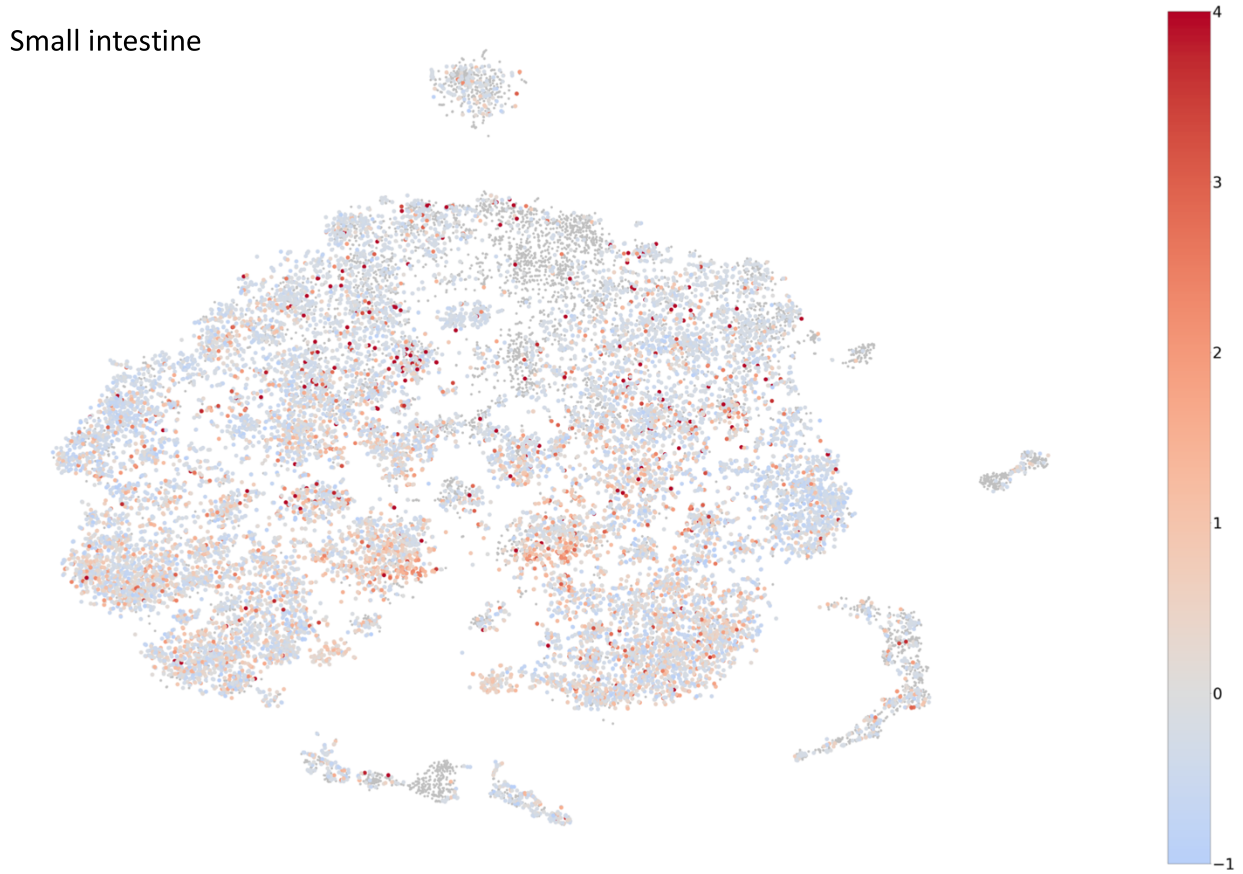


**Figure S24**: Gene co-expression map with genes colored by z-scores obtained using GTEx small intestine expression data.

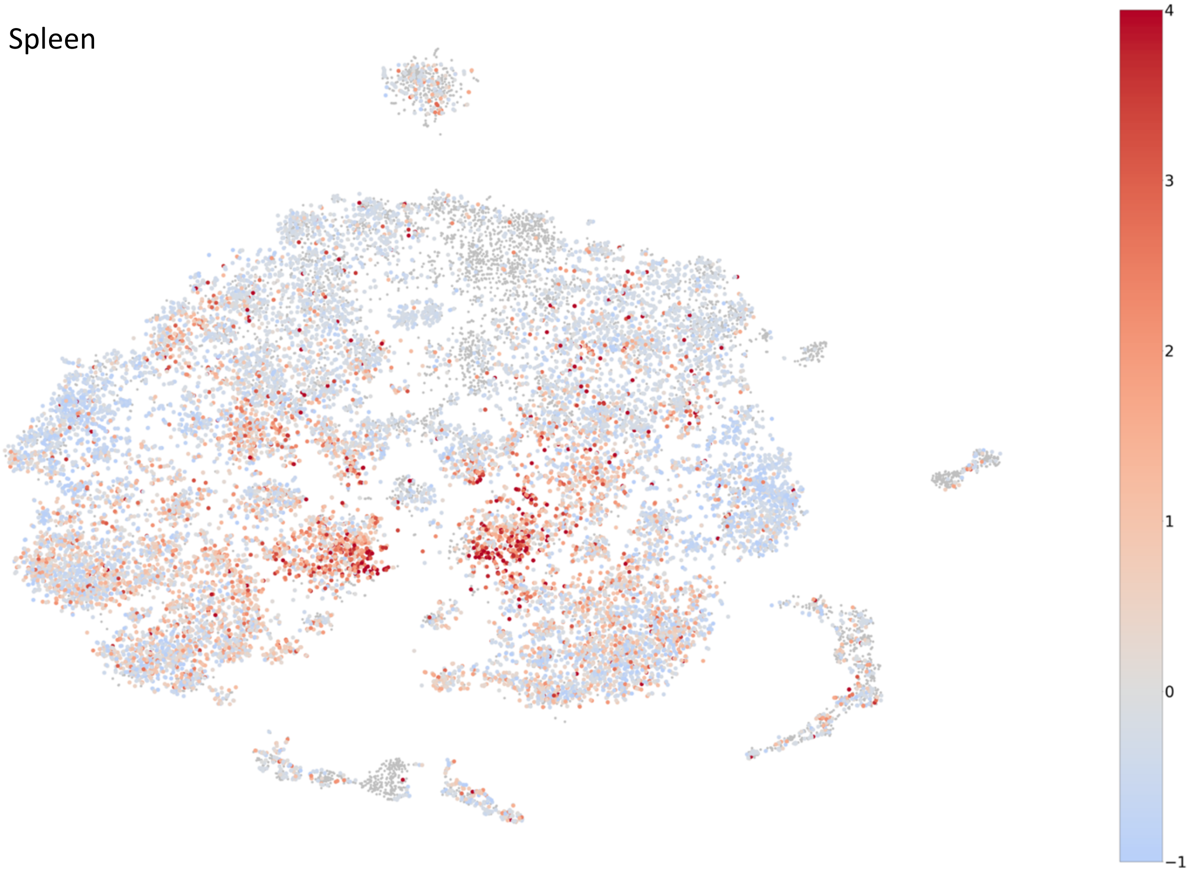


**Figure S25**: Gene co-expression map with genes colored by z-scores obtained using GTEx spleen expression data.

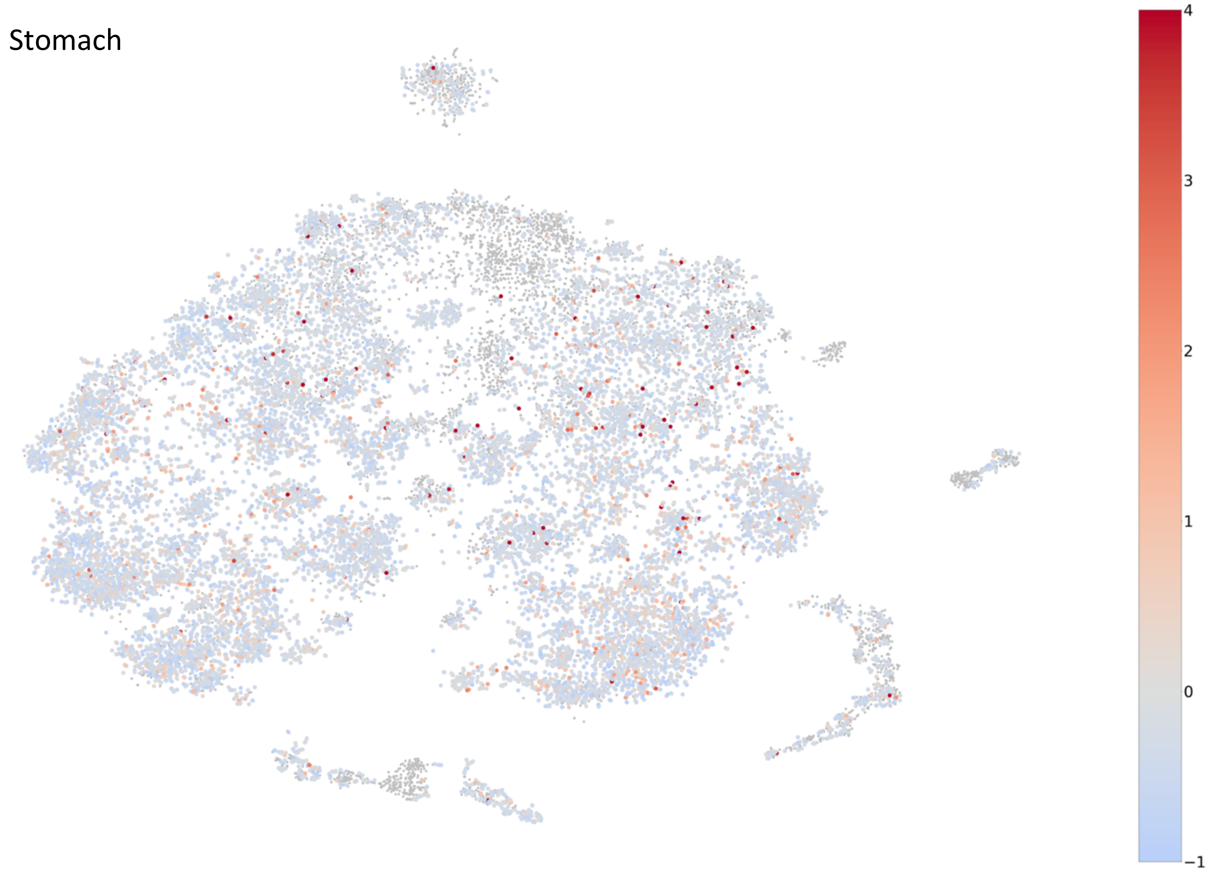


**Figure S26**: Gene co-expression map with genes colored by z-scores obtained using GTEx stomach expression data.

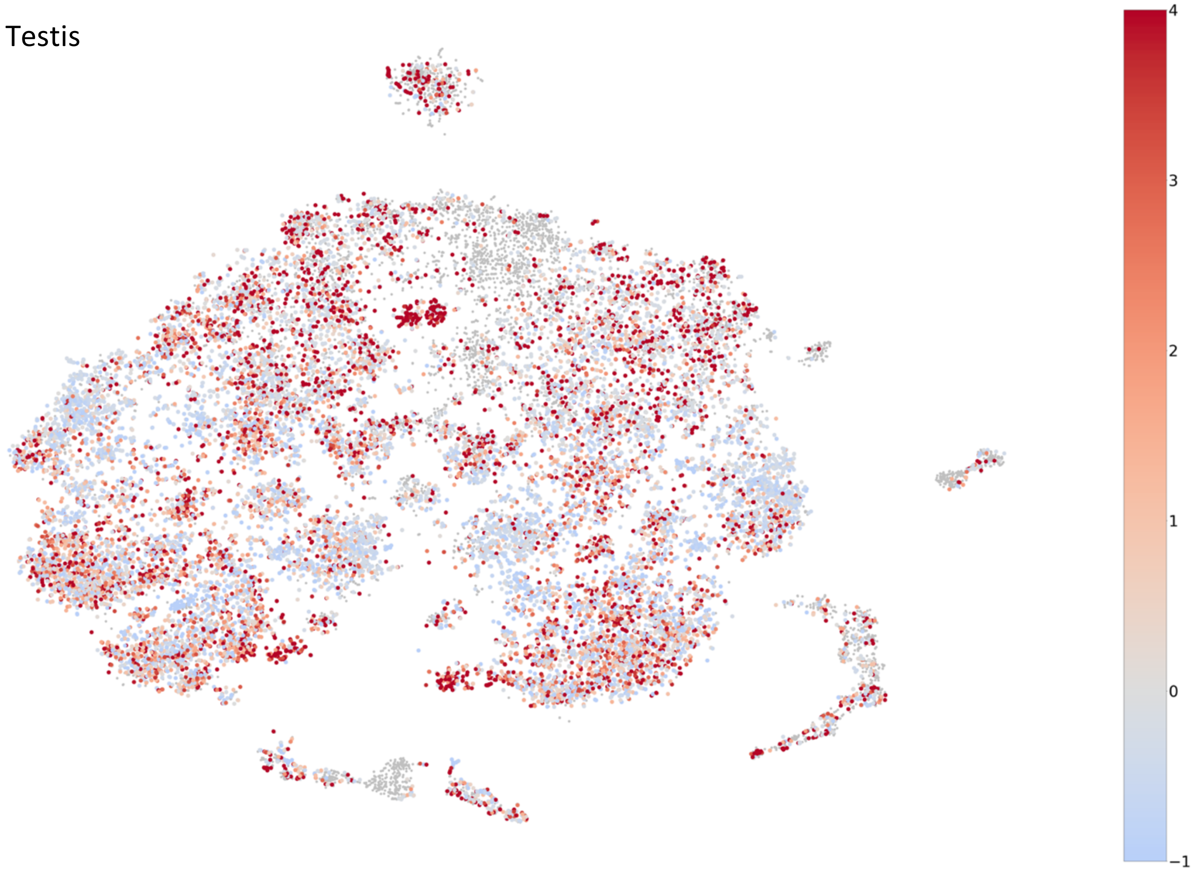


**Figure S27**: Gene co-expression map with genes colored by z-scores obtained using GTEx testis expression data.

Thyroid

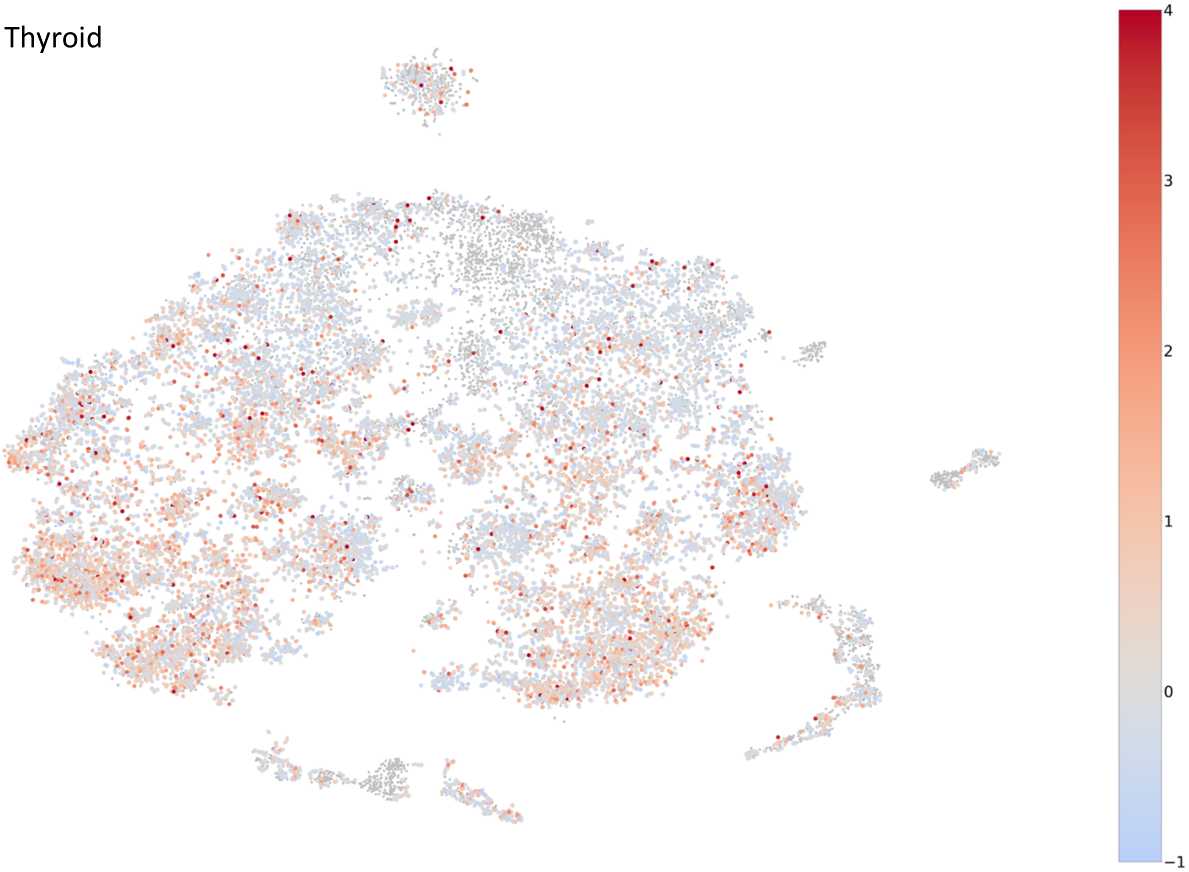


**Figure S28**: Gene co-expression map with genes colored by z-scores obtained using GTEx thyroid expression data.

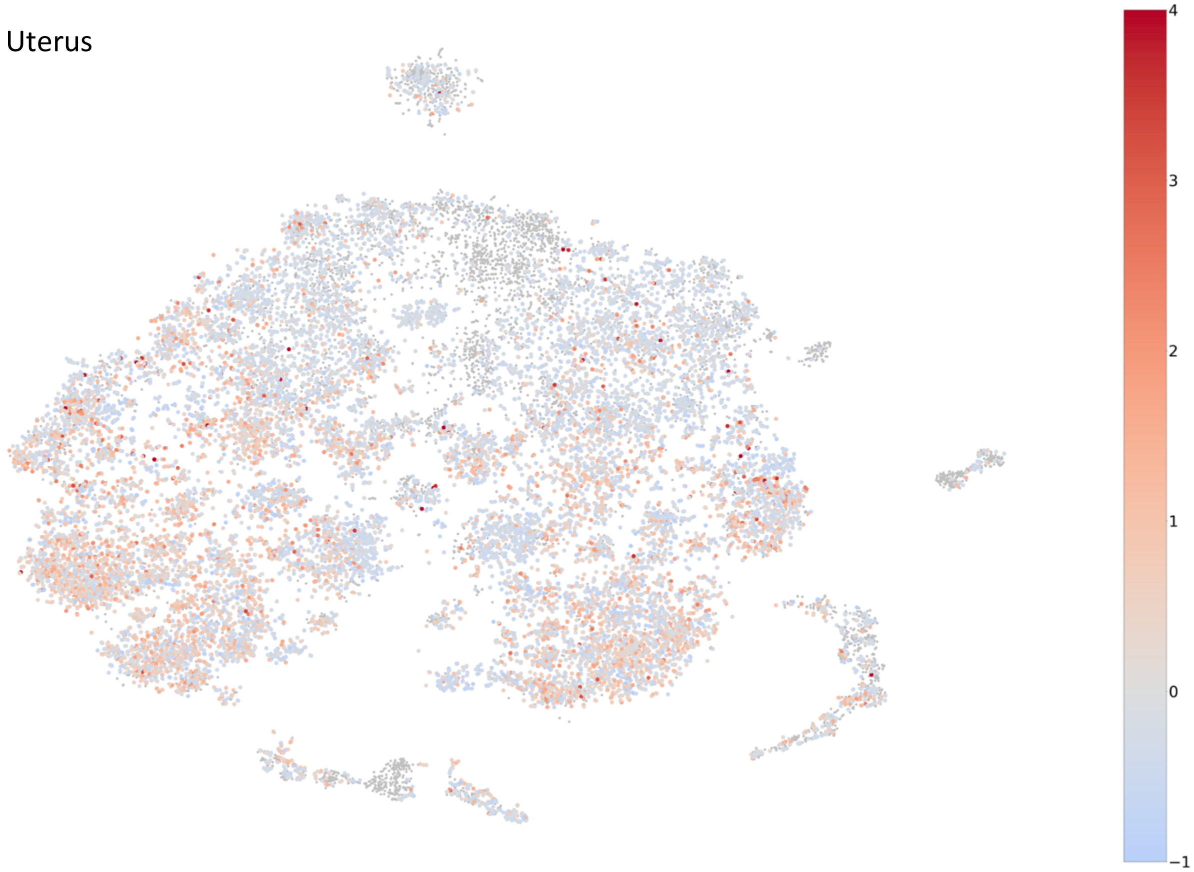


**Figure S29**: Gene co-expression map with genes colored by z-scores obtained using GTEx uterus expression data.

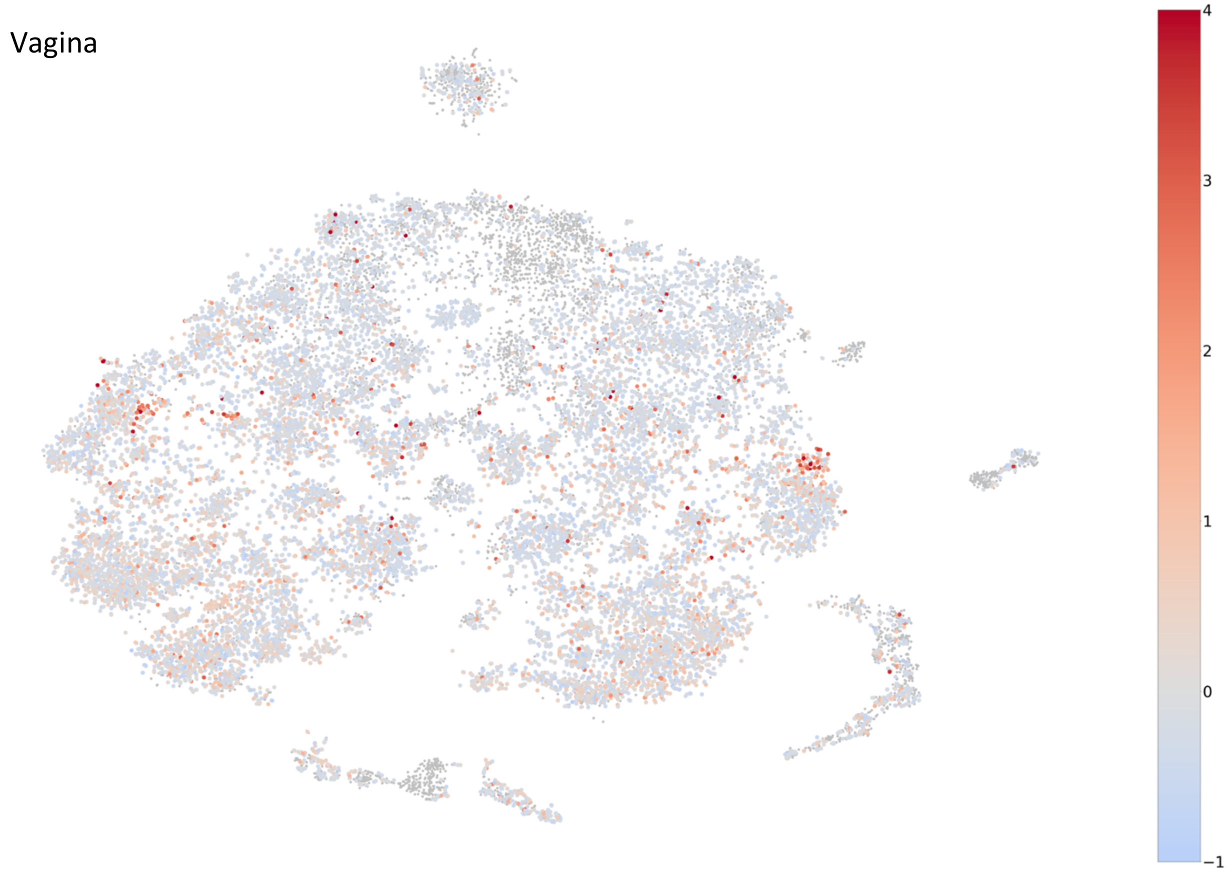


**Figure S30**: Gene co-expression map with genes colored by z-scores obtained using GTEx vagina expression data.
